# Bonsai: An efficient method for inferring large human pedigrees from genotype data

**DOI:** 10.1101/2021.04.06.438656

**Authors:** Ethan M. Jewett, Kimberly F. McManus, William A. Freyman, the 23andMe Research Team, Adam Auton

## Abstract

Pedigree inference from genotype data is a challenging problem, particularly when pedigrees are sparsely sampled and individuals may be distantly related to their closest genotyped relatives. We present a new method that infers small pedigrees of close relatives and then assembles them into larger pedigrees. To assemble large pedigrees, we introduce several new formulas and tools including a new likelihood for the degree separating two small pedigrees, a method for detecting individuals who share background identity-by-descent (IBD) that does not reflect recent common ancestry, and a method for identifying the ancestral branches through which distant relatives are connected. Our method also takes several new approaches that help to improve the accuracy and efficiency of pedigree inference. In particular, we incorporate age information directly into the likelihood rather than using ages only for consistency checks and we employ a heuristic branch-and-bound-like approach to more efficiently explore the space of possible pedigrees. Together, these approaches make it possible to construct large pedigrees that are challenging or intractable for current inference methods. The new method, Bonsai, is available at https://github.com/23andMe/bonsaitree.

## 2. Introduction

The ability to infer complex multi-generational pedigrees from genotype data has many applications ranging from genealogical research to the study of diseases. As human genotyping datasets continue to grow in size, there is increasing interest in computational methods that can reconstruct large pedigrees efficiently and accurately.

Although the problem of pedigree inference has been studied extensively, the majority of pedigree inference methods are designed for non-human species. A major challenge for pedigree reconstruction in non-human populations is that pairwise relationships can be difficult to infer with high accuracy, even when the degree of a relationship is small, because high quality genotype data may be unavailable. As a result, methods typically require that all or most individuals in a pedigree are sampled so that pedigrees can be assembled by connecting strings of parent-child, full-sibling, or half-sibling pairs (Almudevar 2003, Almudevar and Anderson 2012, Cowell 2009; 2013, Cussens et al. 2013, Wang 2004, Jones and Wang 2017, Kirkpatrick et al. 2011, Riester et al. 2009, Sheehan et al. 2014). Although it is possible to connect slightly more distant relationships (Huisman 2017, Anderson and Ng 2016), the majority of existing pedigree inference algorithms can be characterized as methods for either jointly or independently inferring pairwise parent-child pairs and full or half sibling sets, which are then consistent with a pedigree structure when assembled together.

In contrast to non-human pedigrees, genotype data for human populations is comparatively abundant and close relationships, such as parent-offspring or sibling pairs, can be inferred with a high degree of accuracy. The major challenge of pedigree inference in human populations is the fact that pedigrees are often sparsely sampled, with few genotyped sibling and parent pairs and few genotyped individuals beyond the most recent two or three generations. In human datasets, including direct-to-consumer genetic databases, genotyped individuals may have only a few genotyped relatives within a radius extending to first or second cousins and it is common for an individual’s closest relative to be more distant than a second cousin. As a result, it is difficult to construct solid frameworks of close relatives and their genotyped ancestors into which other genotyped individuals can be placed.

There are currently two state-of-the-art methods for inferring complex human pedigrees from genotype data, both of which are maximum likelihood approaches that attempt to find a pedigree that maximizes the sum of log likelihoods of pairwise relationships, given observed patterns of identity-by-descent (IBD) sharing. The two methods differ primarily in the approaches they take to find the maximum likelihood pedigree.

The first and older method, PRIMUS (Staples et al. 2014), explores the space of possible pedigrees by starting with a seed individual and then iteratively adding individuals to the pedigree. Each time an individual is added, the method considers all possible positions that are consistent with the estimated pairwise relationships and the highest likelihood configuration is selected. When adding an individual to the pedigree, each pedigree at the previous step serves as a seed pedigree onto which the individual can be added in multiple ways. By constructing a large set of pedigrees in this way, the algorithm efficiently explores the space of pedigrees that are compatible with the estimated pairwise relationships.

In contrast to PRIMUS, the more recent CLAPPER method (Ko and Nielsen 2017) begins by connecting all individuals together into an initial guess of a pedigree. Then, at each subsequent step, the CLAPPER algorithm rearranges the relationships in the pedigree. This update step is done using a Markov chain Monte Carlo (MCMC) approach in which there are many different possible moves that can be made, such as adding or subtracting a degree of relatedness between two individuals, swapping the labels of two nodes, or pruning off an individual and their descendants and attaching them somewhere else.

The CLAPPER method is typically more accurate than PRIMUS (Ko and Nielsen 2017), whereas the PRIMUS approach can be faster than the MCMC approach used by CLAPPER. However, neither approach was designed to infer the large and sparse pedigrees that are common in direct-to-consumer genetic datasets where the degree of relationship separating a pair of genotyped individuals may be large, verging on degrees where individuals frequently share no detectable IBD. For such pedigrees, searching a broad pedigree space using the approach of PRIMUS or CLAPPER is computationally infeasible. Instead, it is useful to develop an inference approach that dramatically narrows the space of possible pedigrees, while being careful not to exclude the portion of the space containing the true pedigree.

Here, we introduce a new method, Bonsai, for inferring large and sparse pedigrees. To make inference efficient and accurate, we first infer small pedigrees of closely-related individuals using an approach that efficiently explores the space of possible pedigrees. This approach is similar to PRIMUS, but differs in key ways that make the search of the pedigree space both more efficient and more thorough. The small pedigrees are then assembled into larger pedigrees using several new techniques, including a generalized version of the DRUID method of Ramstetter et al. (2018), which allows our method to link distantly related individuals into large and sparsely sampled pedigrees. We refer to the first stage as “Small Bonsai” and to the second stage as “Big Bonsai” (Figure 1). We first describe the small and big Bonsai methods, then use both simulated and real data to investigate the performance of the methods and their components.

**Figure 1.**
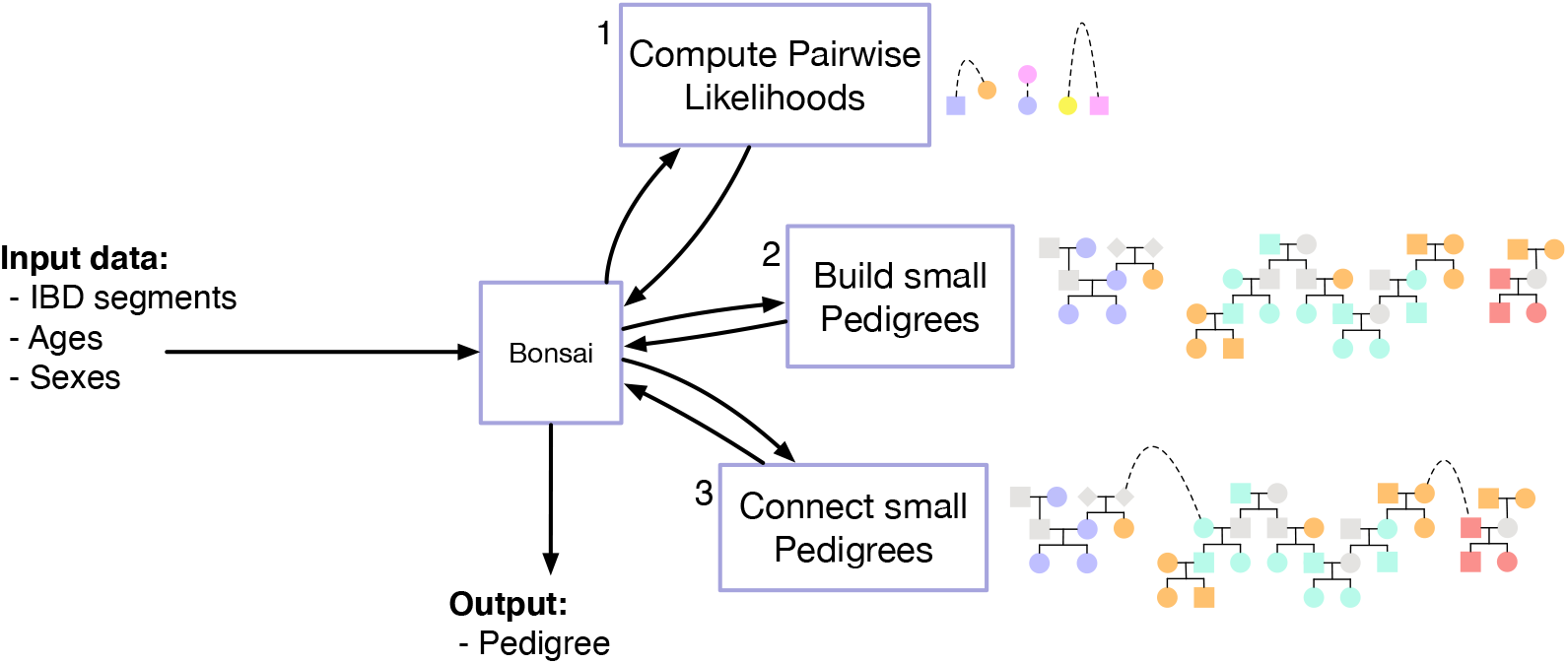
Overview of the full Bonsai method. Details of methods 1, 2, and 3 are presented in Algorithms 1, 2, and 4, respectively.

## 3. Subjects and Methods

### 3.1. Overview of the Bonsai method

The Bonsai method is summarized in Figure 1. The input to the method consists of ages and sexes for a set of putatively-related individuals, along with IBD segments inferred between each pair of individuals. The method then proceeds through three stages in sequence.

First, the relationship between each pair of genotyped individuals is inferred using age and pairwise IBD data. The likelihoods of many other possible relationships are also computed and stored for each pair. Next, small pedigrees of closely-related individuals are inferred from these pairwise likelihoods. Finally, the inferred small pedigrees are assembled into large and sparse pedigrees.

Constructing small pedigrees and combining them together allows us to make use of information in small pedigree structures to improve the accuracy with which more distant relationships are inferred. This approach allows us to more precisely infer the ancestral lineages through which small pedigrees are connected, the number of common ancestors shared by each pair of individuals, and segments of so-called background IBD that do not reflect recent ancestry. Each of these additional pieces of information makes it possible to proactively reduce the space of possible pedigrees that must be searched, making inference tractable for large and sparse pedigrees.

### 3.2. Stage 1: inferring pairwise relationships

The first stage of the Bonsai method is to infer the likelihoods of many possible relationships between each pair of putative relatives. To make the computation of the likelihood efficient without large sacrifices in accuracy, we use a composite likelihood that is the product of the likelihoods of different IBD summary statistics and the likelihoods of the pairwise age differences between the individuals. The genetic component 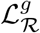 of the likelihood, computed from IBD, is multiplied by the age component 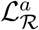 of the likelihood to obtain the final likelihood *ℒ*_*ℛ*_ of a given relationship type, *ℛ*:

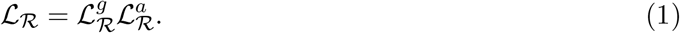

The likelihood is composite, rather than exact, because we do not model the joint distribution of the IBD count and length summary statistics whose product is 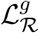 and because there is an underlying joint distribution of IBD sharing and age difference that is not captured by the product of the two likelihoods 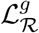 and 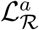.

#### 3.2.1. Pairwise genetic likelihoods

To compute the genetic component of the composite pairwise relationship likelihood, we consider regions of the genome shared identically by descent in a haploid fashion on just one chromatid in each individual, as well as regions shared IBD in a diploid fashion on both chromatids. We use the terms “IBD1 segment” and “IBD2 segment” to refer to regions of haploid and diploid IBD, respectively. The genetic component of the pairwise likelihood is computed using the total length of IBD1 segments, the total length of IBD2 segments, the total number of IBD1 segments, and the total number of IBD2 segments.

It is possible to compute the probability of an observed shared pattern of IBD analytically. However, in practice we find that error in IBD inference leads to differences between the empirical and analytical IBD distributions for each relationship type, especially for close relationships. Thus, we use likelihoods obtained as moment-fitted Poisson and Gaussian approximations of simulated distributions.

Let 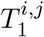 and 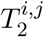 be the total lengths of IBD 1 and 2, respectively for a pair of individuals, *i* and *j* and let 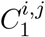 and 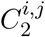 be the counts of the number of IBD 1 and 2 segments shared between two individuals. We follow the convention that uppercase variables *T*_1_, *T*_2_, *C*_1_, *C*_2_, etc. denote random variables and their lowercase counterparts, *t*_1_, *t*_2_, *c*_1_, *c*_2_, etc. denote their observed values. The genetic component of the composite likelihood for a given relationship type, *ℛ*, between a pair of individuals, *i* and *j*, is then computed as

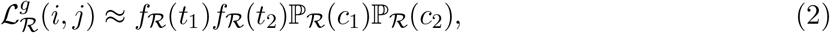

where 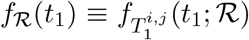 is the probability density function of the sum of lengths of all IBD 1 segments for a relationship of type *ℛ* and ℙ_*ℛ*_ (*c*_1_) *≡*ℙ (*C*_1_ = *c*_1_; *ℛ*) is the probability mass function for the total number of segments of IBD1 for a relationship of type *ℛ*. The quantities *f*_*ℛ*_ (*t*_2_) and ℙ_*ℛ*_ (*c*_2_) are defined analogously for segments of IBD 2.

In Equation (2), the quantities *f*_*ℛ*_ (*t*_1_) and *f*_*ℛ*_ (*t*_2_) are modeled as Gaussian distributions and the distributions ℙ_*ℛ*_ (*c*_1_) and ℙ_*ℛ*_ (*c*_2_) are Poisson with means given by the expected numbers of IBD1 and IBD2 segments, respectively between two individuals of relationship type *ℛ*. The mean and variance of 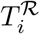, and the mean of 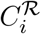 for a relationship of type *ℛ* were obtained empirically using simulations. Details of the simulations used to obtain these moments are provided in Section 3.6.4.

#### 3.2.2. Pairwise age likelihoods

The pairwise age likelihood for a given relationship type, *ℛ*, was obtained by moment-fitting a Gaussian distribution to the differences between the ages of 23andMe customers who self-reported to be of relationship type *ℛ* (Figure 2). We required that the self-reported relationship between each pair of individuals could be verified through a string of inferred parent-child or full-sibling relationships. For example, a self-reported first-cousin relationship between individuals *i* and *j* was verified if *i* and *j* each had inferred parents in the 23andMe database, and if these parents in turn had the same pair of inferred parents, or were inferred to be full siblings.

**Figure 2.**
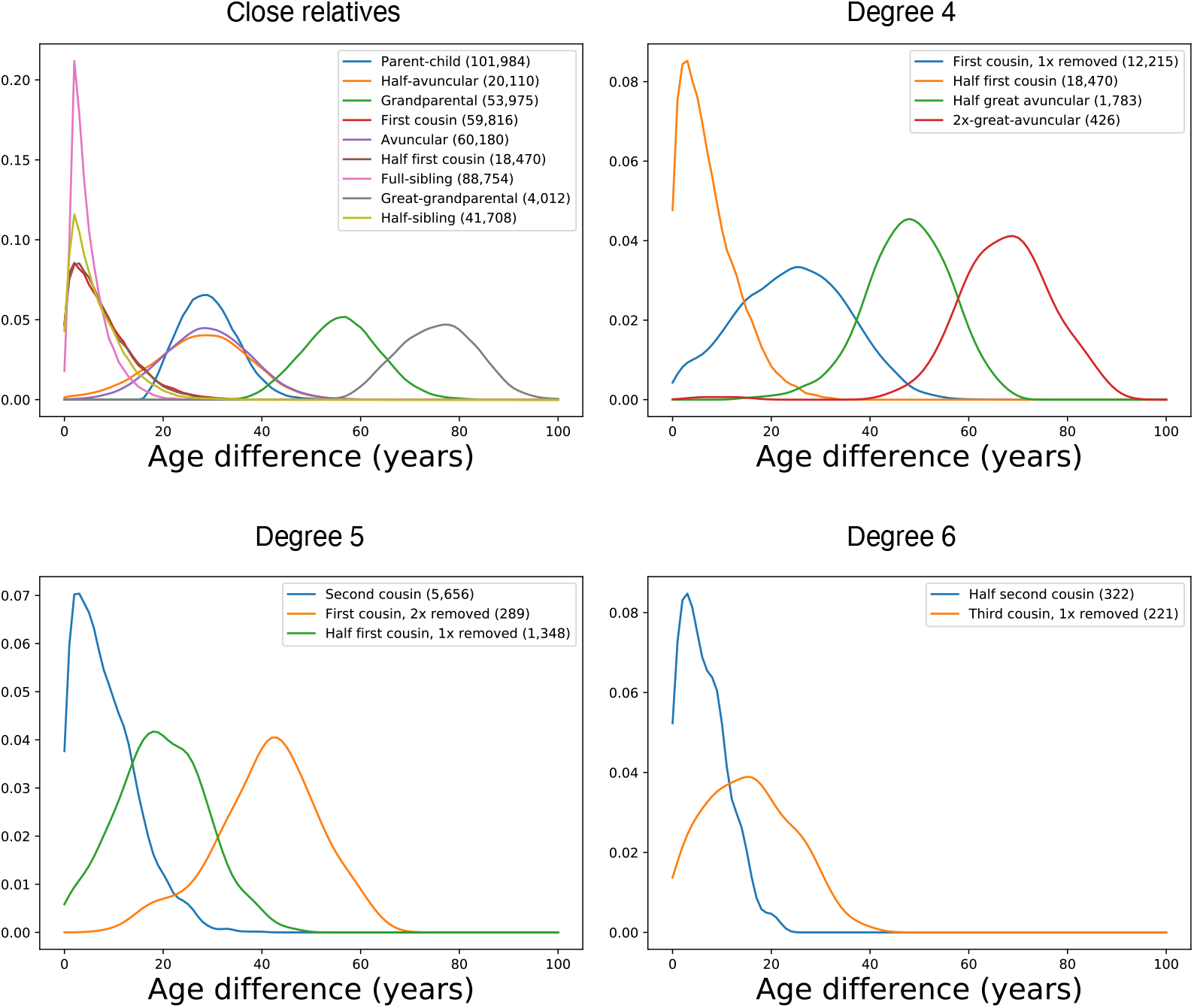
Empirical age difference distributions. Kernel densities for the absolute difference in age between a pair of relatives of a given type. The number of pairs of each type used in the analysis is given in parentheses. Different panels show relationships of different degrees.

For two customers, *i* and *j*, with ages *a*_*i*_ and *a*_*j*_, the age component of the likelihood for relationship type *ℛ* was modeled as a Gaussian distribution with the empirical mean and variance:

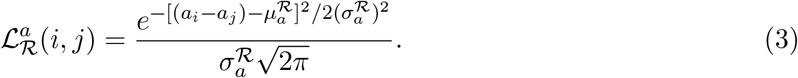

In Equation (3), 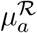 and 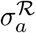 are the moment-fitted mean and standard deviation of the empirical age difference for all pairs of customers who reported the pairwise relationship, *ℛ*. Note that the probability 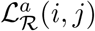 is not symmetrical in the ages, *a*_*i*_ and *a*_*j*_. This is useful for determining the directionality of the relationship between two people, such as parent-child or nephew-aunt when age information is available.

### 3.3. The likelihood of a pedigree

The composite likelihood, *ℒ*_*𝒫*_, of a pedigree *𝒫* is computed as the product of genetic and age likelihoods (Equation 1) for all pairs of individuals in the pedigree,

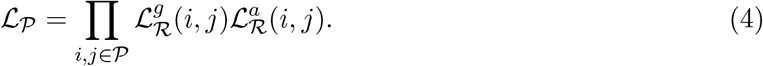

where *ℛ* is the relationship between *i* and *j* implied by the pedigree structure. This likelihood is efficiently computed as each new individual is added to the pedigree. By doing so, we can inductively extend the existing relationships of the parents and/or children of the newly-added person to obtain the relationships of the new person to all existing individuals in the pedigree. We then add the log likelihoods of each of these new pairwise relationships to the log likelihood of the pedigree without the new individual.

### 3.4. The “Small” Bonsai method

To construct a pedigree from pairwise likelihoods, the Small Bonsai method begins by placing a focal individual by itself in the pedigree (Figure 3). This focal individual is typically the person with the closest average degree of relationship to all other individuals in the putatively-related set, but any individual can be chosen. At each subsequent step of the Small Bonsai algorithm, the next individual to be placed is chosen to be the unplaced individual with the closest inferred degree of relationship with one of the individuals already placed in the pedigree, where ties are broken by the total amount of IBD shared. Because each pair of individuals has many possible relationships, we determine the order in which individuals are added using the most likely pairwise relationship for each pair.

**Figure 3.**
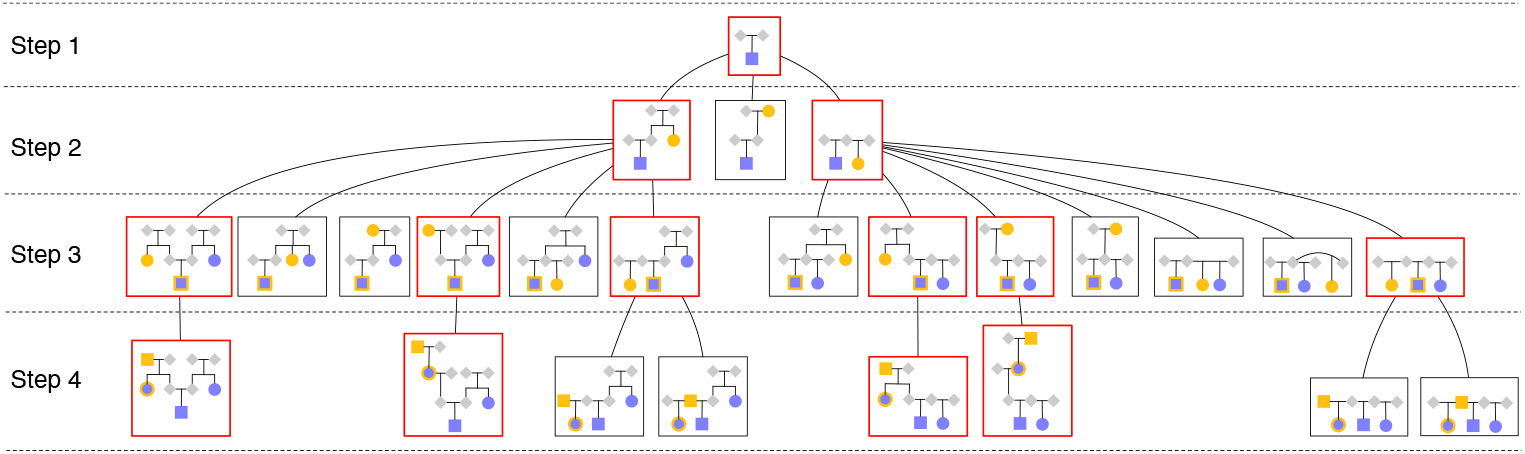
The Small Bonsai method. An example of the sequence of steps for building a small pedigree is shown. The sequence proceeds from top to bottom in the figure. The *i*th row of pedigrees in rectangles represents the *i*th step of the Small Bonsai algorithm in which the *i*th individual is added to a pedigree. Red boxes indicate pedigrees that are retained and carried forward to the next step. Black boxes indicate pedigrees with low likelihoods that are discarded.

The next individual to be placed is considered in all ways that are consistent with the most likely inferred pairwise relationships to individuals already placed. In particular, for a user-specified parameter *r*, we consider the top *r* most likely pairwise relationships between the new individual and their closest relative in the set of placed individuals and we place the individual in all ways that are compatible with each of these *r* most-likely relationships. When adding an individual to the pedigree, we must not only add them in all possible ways to a particular pedigree, we also add them in all *r* ways to all high-likelihood pedigrees that were formed at the previous step.

When two or more pedigrees formed by adding an individual would be topologically identical, we only construct one of the pedigrees. For example, in the second row of Figure 3, because the sexes of the parents are unknown and there are no placed relatives except the focal individual that can be used for triangulation, adding an avuncular relative through the right parent is topologically identical to adding them through the left parent. Therefore, we only build one of these pedigrees (the one on the far left of the second row).

To avoid a rapid expansion in the number of pedigrees at each step, we employ a heuristic branch-and-bound-like procedure in which we discard each pedigree at the end of each step that is very unlikely, compared with the most likely pedigree. In particular, we discard all pedigrees whose likelihoods are less than a fraction *f*_*𝓁*_ of the likelihood of the most likely pedigree, where a pedigree’s likelihood is the product over the likelihoods of the pairwise relationships implied by the pedigree (Section 3.3). In practice, when individuals are closely related, there are only a few pedigrees that have high likelihoods and the rest can be discarded. As a result, the likelihood threshold has a low impact on accuracy while serving to dramatically speed up pedigree building.

This heuristic branch-and-branch-and-bound-likebound-like procedure is repeated until no unplaced individual has a pairwise point-estimated degree that is within a user-specified degree *d* of any placed individual. At this point, the Small Bonsai algorithm is terminated. If unplaced individuals remain, a new focal individual is chosen from among the unplaced individuals and the Small Bonsai algorithm is applied again. The Small Bonsai algorithm is applied repeatedly, choosing a new focal individual each time, until all individuals have been placed into some pedigree.

Figure 3 shows an example sequence for constructing a pedigree using the Small Bonsai method. In the first row of the figure, a focal individual (shaded yellow square) is placed into a pedigree on their own. Grey diamonds indicate their parents, whose sexes are unspecified. In the second row, the unplaced individual with the closest degree of relationship to the placed individual, is placed into the pedigree (yellow circle). The new individual is placed in all ways that are consistent with the top *r* most likely relationships inferred in the pairwise relationship inference step (Section 3.2). Here, we have chosen *r* = 3. These *r* = 3 most likely relationships happen to be “avuncular,” “grandparental,” and “half-sibling” in the example shown. This is the “branch” step of the heuristic branch-and-bound-like procedure.

Before placing the next individual, we evaluate the likelihood of each pedigree, computed as the product of pairwise likelihoods of the relationships induced by the pedigree. We retain only those pedigrees whose likelihoods are at least a fraction *f*_*𝓁*_ of the likelihood of the most likely pedigree. This is the “bound” step of the heuristic branch-and-bound-like procedure.

In the third row of the diagram, the unplaced individual (yellow circle) with the closest degree of relationship to a placed individual is added to all pedigrees that were carried forward from the previous step. The new individual is added to each pedigree in all ways that are consistent with the top *r* most likely relationships to their closest placed relative (purple square with a yellow boundary). Again, these relationships happen to be “avuncular,” “grandparent,” and “half-sibling” in the example. We then perform the “bound” step, retaining only those pedigrees whose likelihoods are at least a fraction *f*_*𝓁*_ of the likelihood of the most likely pedigree.

In the fourth row, we show one final iteration of the procedure. Again, the unplaced individual (yellow circle) is added in all ways that are consistent with the top *r* most likely pairwise point estimated relationships with their closest relative (purple circle with yellow a boundary). In this case the most likely point-estimated relationship happens to be “parental.” Because parent-child relationships are inferred with near certainty, we have only placed the individual as a parent in the diagram, omitting the next 2 most likely relationships which will be considerably less likely.

### 3.5. The “Big” Bonsai method

#### 3.5.1. Overview of the Big Bonsai method

When building a pedigree containing distantly-related individuals, the Small Bonsai method is first applied repeatedly to build sets of small non-overlapping pedigrees. The union of individuals in these small pedigrees is equal to the set of individuals in the full pedigree. The Big Bonsai method is then applied to combine the small pedigrees together, one pair at a time, with the two pedigrees sharing the most total IBD combined at each step.

The Big Bonsai method relies on several new methods we introduce that are useful for different aspects of combining pedigrees together. The first method is a generalized version of the DRUID estimator (Ramstetter et al. 2018) for inferring the degree of relatedness separating the common ancestors of two small pedigrees. The DRUID estimator was derived for specific pedigree structures, such as a set of siblings and their avuncular relatives connected to another such pedigree through the common grandparental ancestors of the two pedigrees. Here, we generalize the DRUID estimator to any pair of outbred pedigrees and, in Appendix 6.3, we further generalize the DRUID estimator to the case in which two pedigrees are connected through two individuals who are not the common ancestors of their respective pedigrees.

The second tool we introduce is an approximation of the likelihood of the degree separating two pedigrees, as a function of the total IBD shared between the two pedigrees. This likelihood, which was inspired by the DRUID estimator, makes it possible to evaluate the relative likelihoods of different degrees separating two pedigrees in addition to obtaining a point estimate of the degree.

The third tool we introduce is a new test for detecting segments of background IBD. Background IBD segments are regions of the genome that are shared identically-by-state (IBS) and which did not arise by transmission from a single shared common ancestor. Instead, these segments arose because of demographic or evolutionary processes, such as a population bottleneck. They are long regions of IBS with hidden recombination events and they can provide misleading information about the degree of relationship between a pair of individuals. Background IBD segments can lead to mis-inferred pedigrees, particularly when pedigrees are sparsely genotyped.

**Table 1.**
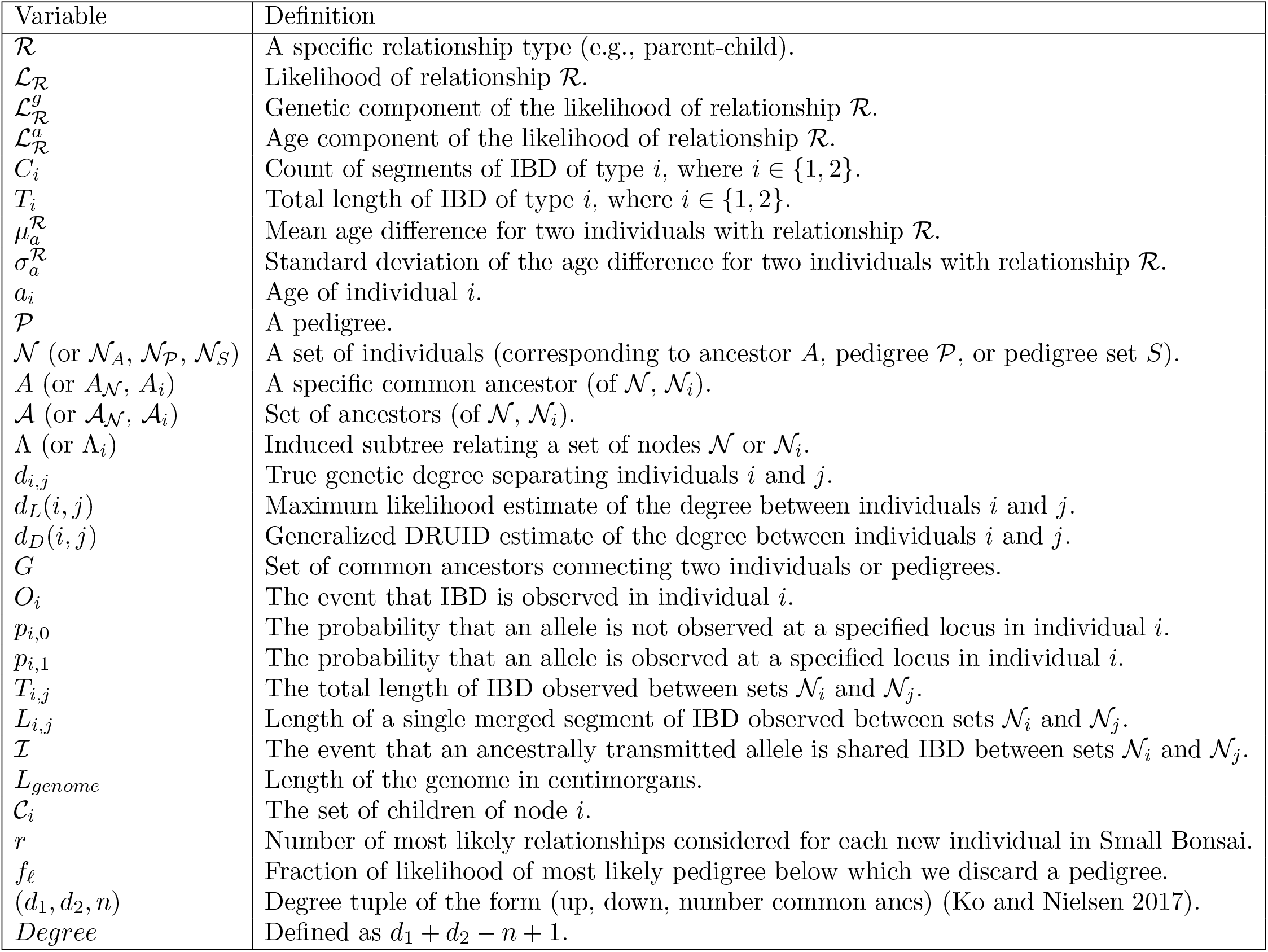
Variable definitions.

The fourth tool we introduce is a method for determining the correct ancestral lineages through which two or more pedigrees are connected. This approach relies on detecting overlapping IBD segments that are inconsistent with certain lineage combinations.

We also derive a recursive formula for computing the probability of an observed presence-absence pattern of an ancestrally transmitted allele in their descendants. This formula is useful for developing the generalized DRUID estimator and the likelihoods for degree estimation and background IBD detection.

Together, the new tools we introduce can be used to identify the ancestors through which two small pedigrees are connected, infer the degree separating the two ancestors, and identify and discard individuals whose IBD sharing patterns appear to be background IBD. By using these inference tools to identify highly-likely ways of connecting pedigrees, the space of possible pedigrees can be considerably reduced. We now describe each of these approaches in detail.

#### 3.5.2. The probability of a presence-absence pattern of an ancestral allele

Consider two pedigrees *𝒫*_1_ and *𝒫*_2_ of genotyped individuals, *𝒩*_1_ and *𝒩*_2_, related through a common ancestor (or pair of ancestors), *G* (Figure 4). Let *A*_1_ be the common ancestor of *𝒩*_1_ in *𝒫*_1_ and let *A*_2_ be the common ancestor of *𝒩*_2_ in *𝒫*_2_.

**Figure 4.**
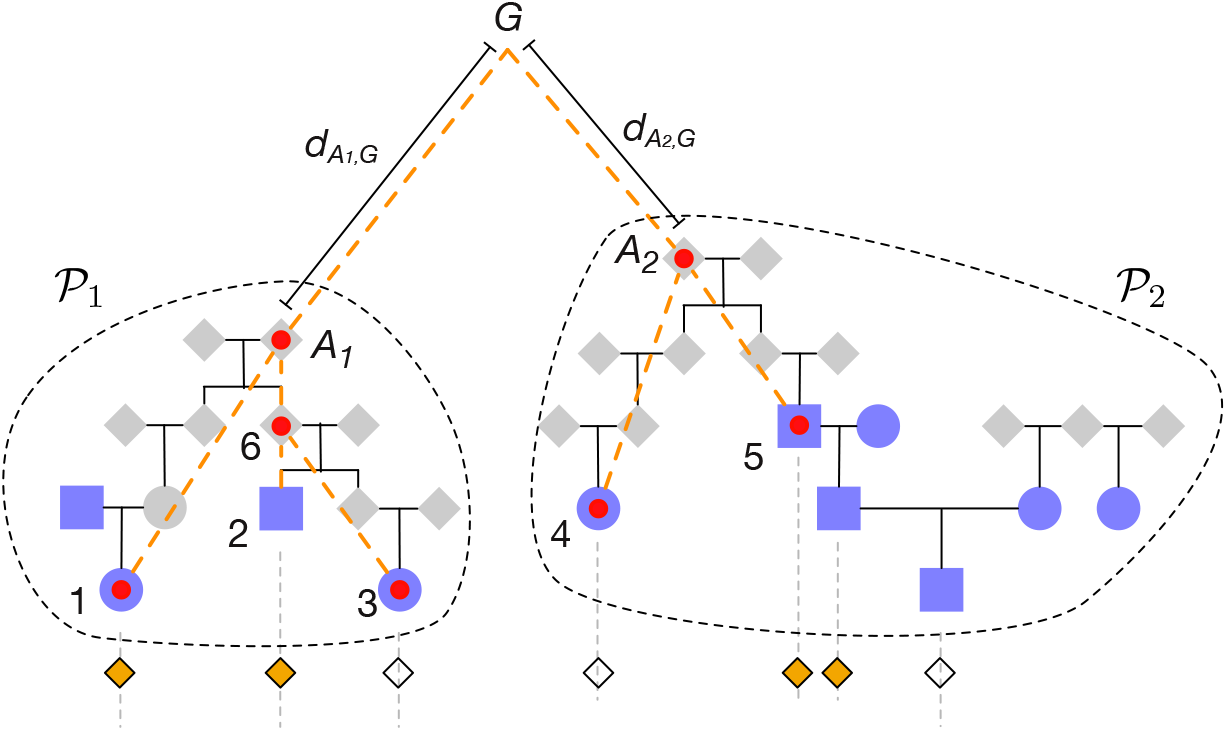
Example of an observed pattern of presence and absence of an ancestral allele. Genotyped individuals are shaded in purple. Filled and empty diamonds below indicate the presence or absence of the allele from *G*. Red dots on purple genotyped individuals indicate the set of genotyped individuals with no direct genotyped ancestors. Red dots on grey ungenotyped individuals indicate the most recent common ancestors transmitting the segments to the genotyped individuals. Dashed orange lines indicate the paths by which the allele is transmitted from common ancestor *G*. The number of meioses separating *A*_1_ and *A*_2_ from a common ancestor, *G*, are 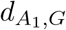 and 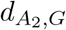.

Consider an allele transmitted from one chromatid in *G* to its descendants. We begin by deriving the probability of the observed pattern of presence and absence of the ancestral allele among descendants of *A*_1_ and *A*_2_. Let 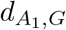 and 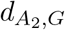 be the degrees separating *A*_1_ and *A*_2_ from the set of most recent common ancestors, *G*, of the pedigree. *G* corresponds to two individuals if *A*_1_ and *A*_2_ are descended from an ancestral couple and *G* corresponds to a single common ancestor if *A*_1_ and *A*_2_ are descended from a pair of half siblings. We do not consider cases of endogamy, where *G* corresponds to more than one ancestor other than a mate pair.

Figure 4 shows a presence-absence pattern of an inherited allele among genotyped individuals in the two small pedigrees *𝒫*_1_ and *𝒫*_2_. The probability of the observed presence and absence pattern can be computed recursively by conditioning on whether the allele was observed in the ancestor of each individual. This approach is similar to Felsenstein’s tree pruning algorithm (Felsenstein 1981).

Let *O*_*i*_ be a random variable describing the event that a copy of the allele is transmitted to descendant *i* and is observed. We set *O*_*i*_ = 1 if the allele is observed in individual *i* and *O*_*i*_ = 0 if it is not observed. The probabilities ℙ (*O*_*i*_ = 0) and ℙ (*O*_*i*_ = 1) can be computed by conditioning on whether the allele in *G* was observed in the node of the induced subtree immediately ancestral to *i*.

Defining

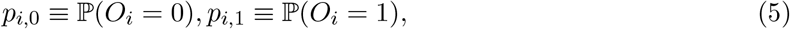

we show in Appendix 6.1 that the probabilities can be computed using the recurision

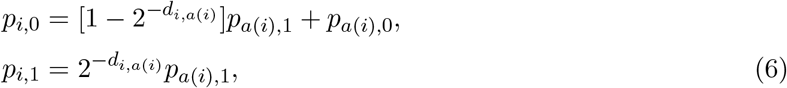

with the base conditions *p*_*g*,0_ = 0 and *p*_*g*,1_ = 1 for each allelic copy, *g*, in *G*. The probability of an observed IBD sharing pattern {*O*_1_, …, *O*_*k*_} across *k* leaf nodes can be computed recursively using Equation (6).

#### 3.5.3. The generalized DRUID estimator

Ramstetter et al. (2018) developed a method for inferring degrees of relatedness among distant relatives. The method addresses the problem that the amount of IBD shared between two individuals decreases exponentially with their degree of relatedness, resulting in very little information for inferring degrees between distant relatives. In fact, there can be a non-negligible probability that distant relatives will share no IBD segments at all, especially if information contained in short IBD segments is discarded to reduce the rate of false positive IBD segments.

Because two genealogically-related individuals may share little or no IBD, it is helpful to leverage IBD segments shared among close relatives of the two individuals when inferring their degree of relatedness. Figure 5 illustrates the utility of considering IBD segments among groups of individuals rather than pairwise IBD when the degree of relatedness is not small. In particular, individuals 3 and 4 in Figure 5 share no IBD segments. Thus, one cannot infer their degree of relatedness without additional information. However, if close relatives of 3 and 4 do share IBD with one another, and if pedigrees can be inferred relating these close relatives to 3 and 4, then we can use the IBD in these relatives to estimate the degree of relationship between 3 and 4.

**Figure 5.**
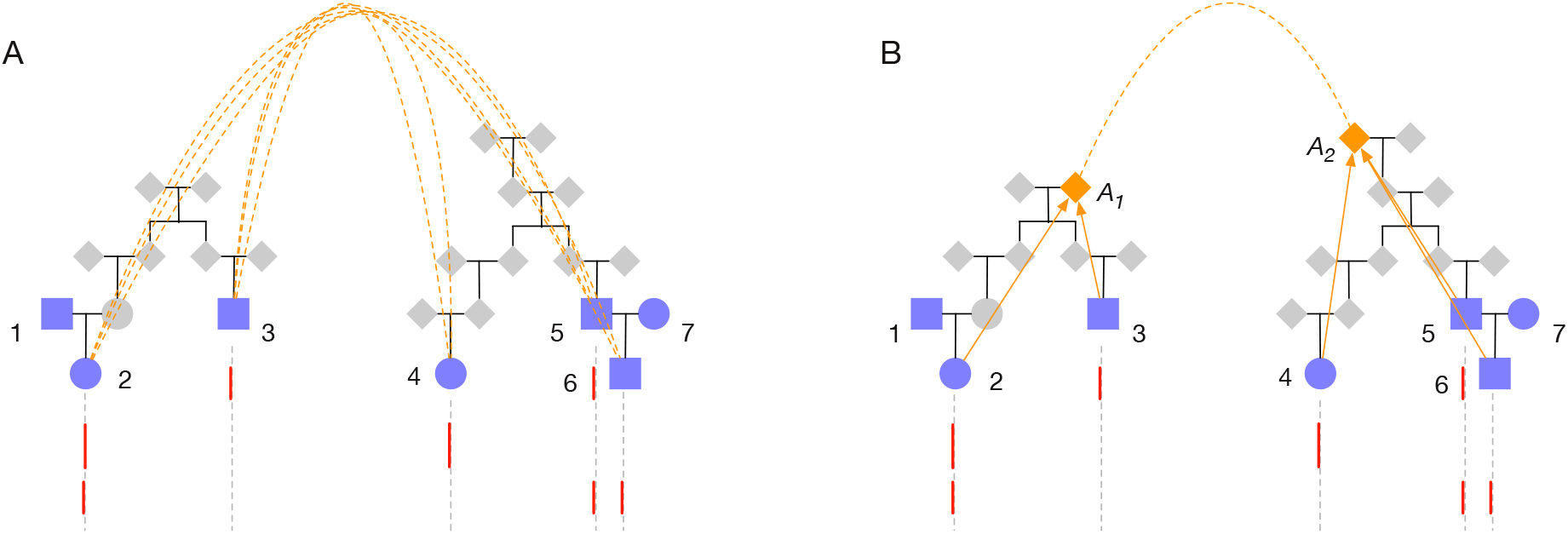
Leveraging IBD from close relatives to infer the degree of relatedness between individuals. Each panel in the figure shows a comparison between two pedigrees. Purple-shading indicates individuals who have been genotyped. Red lines indicate IBD shared between a genotyped individual in the pedigree containing 1, 2, and 3 and a genotyped individual in the pedigree containing 4, 5, 6, and 7. Orange dashed lines in Panel (A) indicate pairwise degrees of relatedness among all cross-pedigree pairs. The orange dashed line in Panel (B) indicates the degree of relatedness between the common ancestor *A*_1_ of the IBD-carrying individuals in the left pedigree and the common ancestor *A*_2_ of the IBD-carrying individuals in the right pedigree. In Panel (A), pairwise IBD is summarized to infer the degree separating the two pedigrees. In Panel (B), IBD information in the descendants of *A*_1_ and *A*_2_ is summarized to infer the degree of relatedness among these two common ancestors. The approach in Panel (A) is taken by the PADRE method (Staples et al. 2016). The approach in Panel (B) is taken by the DRUID method (Ramstetter et al. 2018).

Leveraging IBD shared by close relatives has the effect of increasing the amount of available data for inferring pairwise relationships. Ramstetter et al. (2018) demonstrated that an approach based on summarizing IBD from close relatives can greatly improve the accuracy of estimates of distant degrees of relatedness compared with the approach of computing a composite likelihood over all pairs of individuals (Staples et al. 2016). These two approaches are shown in Figure 5.

Let *𝒩*_1_ and *𝒩*_2_ be two sets of genotyped individuals; for example, sets *𝒩*_1_ = {2, 3} and *𝒩*_2_ = {4, 5, 6} in Figure 5B. Let *A*_1_ and *A*_2_ be the most recent common ancestors of *𝒩*_1_ and *𝒩*_2_, respectively and let *d*(*A*_1_, *A*_2_) denote the degree between *A*_1_ and *A*_2_. The DRUID estimator of *d*(*A*_1_, *A*_2_) derived by Ramstetter et al. is obtained by first merging all IBD segments observed between *𝒩*_1_ and *𝒩*_2_. The total merged IBD is then converted into a point estimate of the amount of IBD shared between the common ancestor *A*_1_ and the common ancestor *A*_2_.

The amount of IBD shared between *A*_1_ and *A*_2_ is estimated by considering the fraction *φ*_1_ of the genome of *A*_1_ that is passed on to its genotyped descendants in *𝒩*_1_ and the fraction *φ*_2_ of the genome of *A*_2_ that is passed on to its genotyped descendants in *𝒩*_2_. If *IBD*(*A*_1_, *A*_2_) is the amount of IBD shared between *A*_1_ and *A*_2_, then the expected amount shared between *𝒩*_1_ and *𝒩*_2_ is *IBD*(*𝒩*_1_, *𝒩*_2_) = *φ*_1_*φ*_2_*IBD*(*A*_1_, *A*_2_). Solving for *IBD*(*A*_1_, *A*_2_) yields a point estimator of *IBD*(*A*_1_, *A*_2_) in terms of the observed quantity *IBD*(*𝒩*_1_, *𝒩*_2_).

Ramstetter et al. (2018) derived formulas for *φ*_1_ and *φ*_2_ for specific pedigree configurations, such as sets of siblings or siblings together with avuncular relatives. Here, we generalize the DRUID estimator to arbitrary outbred pedigrees and further generalize the method to include the case in which *A*_1_ is descended from a descendant *A*_2_ who is ancestral to only a subset of genotyped individuals in *𝒩*_2_.

The fraction *φ*_*i*_ of the genome of *A*_*i*_ that is passed on to some descendant in *𝒩*_*i*_ can be computed as

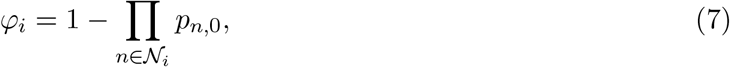

where the quantities *p*_*n*,0_ are computed recursively using Equation (6). Thus, an estimate of the amount of IBD shared between *A*_1_ and *A*_2_ is

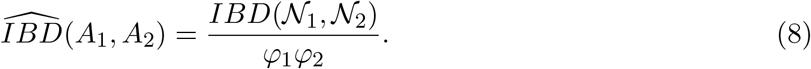

Using the expression 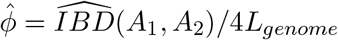 for the kinship coefficient when all IBD is of type 1, we obtain the generalized DRUID estimator

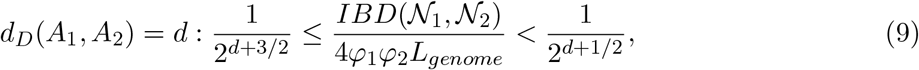

where the bounds come from Manichaikul et al. (2010) and are the ones used for the DRUID estimator presented in Ramstetter et al. (2018).

In Appendix 6.3, we demonstrate how the DRUID estimator can be further generalized to the case in which *A*_1_ is descended from one of the individuals in *𝒩*_2_, or from an internal node of the induced subtree that is a descendant of *A*_2_. Thus, we obtain a version of the DRUID estimator that can be applied to general outbred pedigrees.

#### 3.5.4. The likelihood of the degree of relatedness among groups of individuals

Using the DRUID principle, we can develop a likelihood estimator of the pairwise degree of relatedness between the common ancestors *A*_1_ and *A*_2_, given the observed total IBD *T*_1,2_ between the genotyped descendants of *A*_1_ and *A*_2_.

Consider again the scenario depicted in Figure 4 in which two sets of genotyped individuals, *𝒩*_1_ and *𝒩*_2_, are related through a common ancestor or pair of ancestors, *G*. The probability that a given allele from *G* is observed IBD between *𝒩*_1_ and *𝒩*_2_ can be obtained by conditioning on the events that it is observed in *A*_1_ and *A*_2_. Let *ℐ* denote the event that the allele is observed IBD. Then

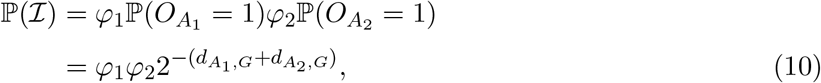

where *φ*_*i*_ is computed using Equation (7).

If *A*_1_ and *A*_2_ had exactly one common ancestor with one allele to transmit, then Equation (10) would be the fraction of the genome in which we expect to find some segment shared IBD between some member of *𝒩*_1_ and some member of *𝒩*_2_. However, we must now account for the fact that each common ancestor of *A*_1_ and *A*_2_ in *G* carries two allelic copies and that there can be either one or two such common ancestors.

Let |*G*| denote the number of common ancestors of *A*_1_ and *A*_2_, each of which carries two alleles at the locus of interest. Let *ℐ* ^*c*^ denote the complement of event *ℐ*, i.e., the event that a specific allele from *G* is not observed IBD. Thus, we have

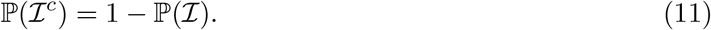

Then the probability that none of the 2|*G*| alleles is observed IBD is ℙ (*ℐ* ^*c*^)^2|*G*|^, and the probability that at least one of the alleles is observed is 1 *−* ℙ (*ℐ* ^*c*^)^2|*G*|^.

We can use the probability of observing an allele IBD to obtain an approximate likelihood of the total length *T*_1,2_ of IBD observed between descendants of *A*_1_ and *A*_2_. The mean of this distribution is simply the expected length of the genome in a state of IBD between the two clades, which is

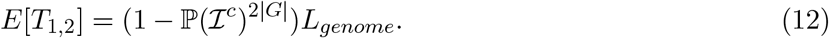

An approximation of the variance of *T*_1,2_ is derived in Section 6.2 and is given by

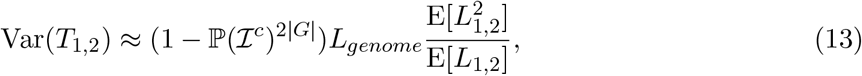

where *L*_1,2_ is the length of any given IBD segment between *A*_1_ and *A*_2_ formed by merging all IBD segments between leaf nodes in *A*_1_ and *A*_2_ that overlap one another. The moments 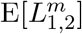 are derived in Appendix 6.2 and can be computed using Equation (28) or (29).

If the segments, *L*_1,2_ were each exponentially distributed, then *T*_1,2_ would have a gamma distribution. In practice, a gamma distribution is an accurate approximation for the distribution of *T*_1,2_, given that the length *T*_1,2_ is greater than zero. Thus, we can approximate the distribution of *T*_1,2_ by

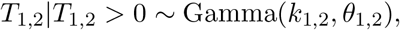

where *k*_1,2_ and *θ*_1,2_ are found by matching the mean and variance of the gamma distribution with E[*T*_1,2_] and Var(*T*_1,2_). Thus, we obtain

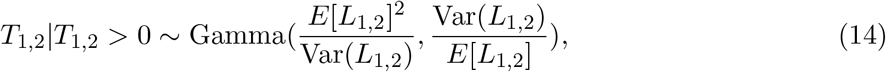

where E[*L*_1,2_] and 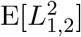 are given by Equation (29).

If every IBD segment has some length, we can assume that *T*_1,2_ is only identically zero when there are no IBD segments. The distribution of the number of segments can be modeled as a Poisson random variable with mean E[*N*_1,2_] equal to the expected number *N*_1,2_ of merged segments shared between *𝒩*_1_ and *𝒩*_2_. The probability that there are no segments is then 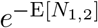. Thus, we have the approximation

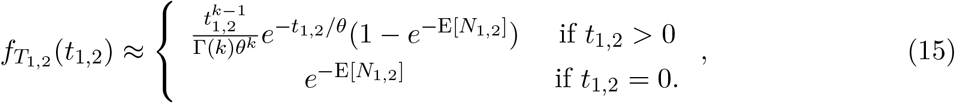

where *k* = *E*[*L*_1,2_]^2^*/*Var(*L*_1,2_), *θ* = Var(*L*_1,2_)*/E*[*L*_1,2_] and E[*N*_1,2_] is given in Equation (25). Figure S2 shows analytical values computed using Equations (12) and (13) compared to empirical values from simulations. Figure S3 shows the approximate analytical distribution computed using Equation (15) compared to the empirical distribution computed from simulations. Although the gamma distribution in Equation (15) provides a good fit to the empirical distribution, a Gaussian distribution can be more robust in practice because the gamma approximation is slightly underdispersed compared with the true distribution. In practice, we use the Gaussian distribution for inference.

A maximum likelihood estimator of the degree between *A*_1_ and *A*_2_ can be obtained by determining the degree *d*_*L*_(*A*_1_, *A*_2_) between *A*_1_ and *A*_2_ for which value of the distribution in Equation (15) is maximized. This gives the maximum likelihood estimator

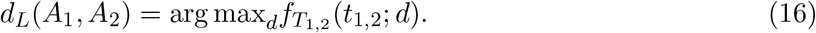

#### 3.5.5. Likelihoods for identifying background IBD

Individuals with no recent relationship can share small segments of IBD by chance, especially in populations with recent or severe bottlenecks. This kind of IBD is referred to as background IBD and it poses a considerable challenge to accurate pedigree inference.

Previous methods have addressed background IBD by various approaches. For example, the authors of the ERSA method (Huff et al. 2011) presented an approach for modeling the distribution of background IBD among unrelated individuals and then performing a likelihood ratio test to determine whether the IBD shared between a new pair of individuals was significantly different from background.

Power for detecting background IBD can be increased by comparing sets of individuals rather than pairs of individuals, leveraging the information inherent in previously-inferred pedigree structures. As we demonstrate, such an approach makes it possible to detect background IBD between sets of individuals without prior knowledge of the distribution of background IBD. This is useful because it can be challenging to know a priori the expected amount of background IBD between a given pair of individuals.

We take an approach to identifying background IBD in which we consider the information contained in IBD sharing patterns across multiple individuals to determine when IBD is background and when it is due to true recent ancestry. In particular, we consider the problem in which all of the IBD observed in an individual is either background IBD, or true IBD due to a recent relationship.

To illustrate the approach, consider the IBD sharing pattern shown in Figure 6. Individuals 3 and 4 share relatively large amounts of IBD with 5 and 6, compared with the amount shared between {1, 2} and {5, 6}. If 1 and 2 were much more distantly related to 5 and 6 than 3 and 4, we might not consider the amount of IBD they share with 5 and 6 to be unusually small. However, because 1, 2, 3, and 4 have similar degrees of relatedness to 5 and 6, the amount of IBD shared by 1 and 2 appears to be unusually low. If we can say that the amount of IBD shared below node 7 is smaller than expected by chance, then we can assume that the IBD observed in 1 and 2 is background IBD and remove these nodes from consideration when connecting the left and right pedigrees.

**Figure 6.**
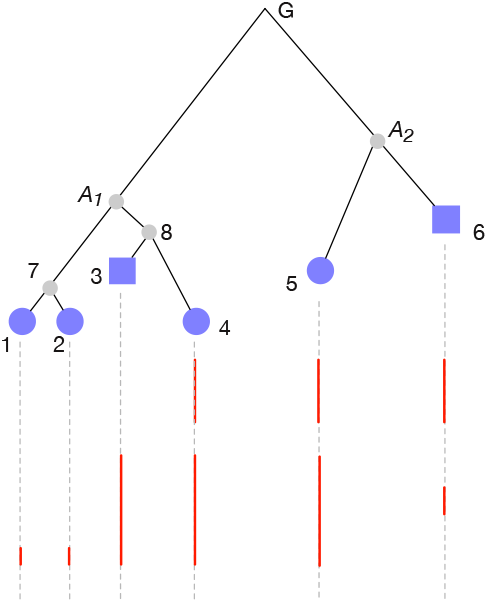
Detecting background IBD. Genotyped individuals are shaded in purple. Vertical red lines indicate IBD segments shared between the genotyped descendants of *A*_1_ and the genotyped descendants of *A*_2_.

We test for background IBD in practice through a series of hypothesis tests. Given that IBD is observed between two sets of nodes, *𝒩*_1_ and *𝒩*_2_, we take the putative common ancestors *A*_1_ and *A*_2_ through which the IBD was inherited to be the most recent common ancestors of *𝒩*_1_ and *𝒩*_2_, respectively. We then consider each of the descendant nodes, *c*, immediately below *A*_1_ in turn (e.g., 7 and 8 in Figure 6) and we ask whether the amount of observed IBD below the node is much lower or higher than expected by chance, given the degree between *A*_1_ and *A*_2_ inferred using all the descendant nodes below *A*_1_, excluding *c*.

If we assume that some individuals in *𝒩*_1_ are related to some individuals in *𝒩*_2_, then on average the observed IBD will represent true IBD, plus background IBD. The individuals sharing the greatest amount of IBD, relative to their genealogical positions, are likely to be the truly-related individuals. When testing for background IBD, we assume that the individuals sharing the greatest amount of IBD are truly related and we test for background IBD only in the individuals sharing less IBD. Thus, the child node *c*^*∗*^ of *A*_1_ with the greatest IBD sharing with *𝒩*_2_ is exempt from our test.

We drop all nodes that reject the null hypothesis of this test and re-set the ancestral node to be the common ancestor of all remaining IBD-carrying nodes. For example, if we detected that the clade below node 7 in Figure 6 had much lower IBD than expected by chance, we would drop node 7 and its descendants from consideration and set the true common ancestor relating the two pedigrees to be node 8. We iteratively repeat this procedure until no nodes are dropped. We then repeat the procedure for the nodes immediately below *A*_2_.

Let *𝒞*_*n*_ denote the set of children of node *n*. To test whether the IBD observed below a child node *𝒸 ∈𝒞*_*n*_ is background IBD, we establish an approximation of the null hypothesis *H*_0_ that the observed IBD below node *𝒸* is real and we ask whether this hypothesis is rejected in favor of the alternative hypothesis *H*_1_ that the IBD is background.

Under *H*_0_, we assume that the degree 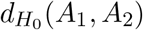 between *A*_1_ and *A*_2_ is the maximum likelihood estimate 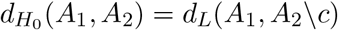, or the generalized DRUID estimate 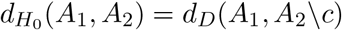 ignoring clade *𝒸*. We then perform the following test

Reject *H*_0_ at level *α* if:

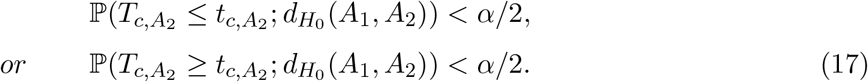

where 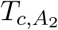 is the random variable describing the amount of IBD between descendants of *c* and descendants of *A*_2_ with observed value 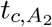. The distribution of 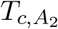 is given by Equation (15). It is reasonable to be conservative when dropping background IBD so that true relationships are called as background IBD only a small fraction of the time. Thus, in practice, we take *α* to be small, such as *α* = 10^*−*3^.

#### 3.5.6. Determining the ancestral branches through which to connect pedigrees

One difficulty in constructing large pedigrees is determining the ancestors through which two sets of gentoyped individuals are related. A simple fundamental question is whether two lineages are both on the maternal side of an individual, both on the paternal side, or on opposite parental sides. Without genotyped parents, the side through which a lineage passes can be difficult to determine, although sex chromosomes and mitochondrial haplotypes can be used to resolve the parent of origin in some cases.

We consider the problem of inferring whether two distant sets of relatives are related through the same parent of a focal individual, or through different parents. The scenario we consider is illustrated in Figure 7. The amount of IBD shared among the red and purple pedigrees in Figure 7 is uninformative about whether they are related through the same parent. Even if the purple and red pedigrees in Figure 7 shared no IBD, they could still be related to individual 1 through the same parent by passing through different grandparents. However, if the red and purple pedigrees are related to the focal individual 1 through the same parent, the IBD segments the purple pedigree shares with individual 1 cannot spatially overlap with the segments the red pedigree shares with individual 1. This is because two overlapping segments would have undergone recombination in the parent (i.e., individual 10). The result will either be a spliced segment (Figure 7), or the replacement of one segment by the other with possible reduction in segment size.

**Figure 7.**
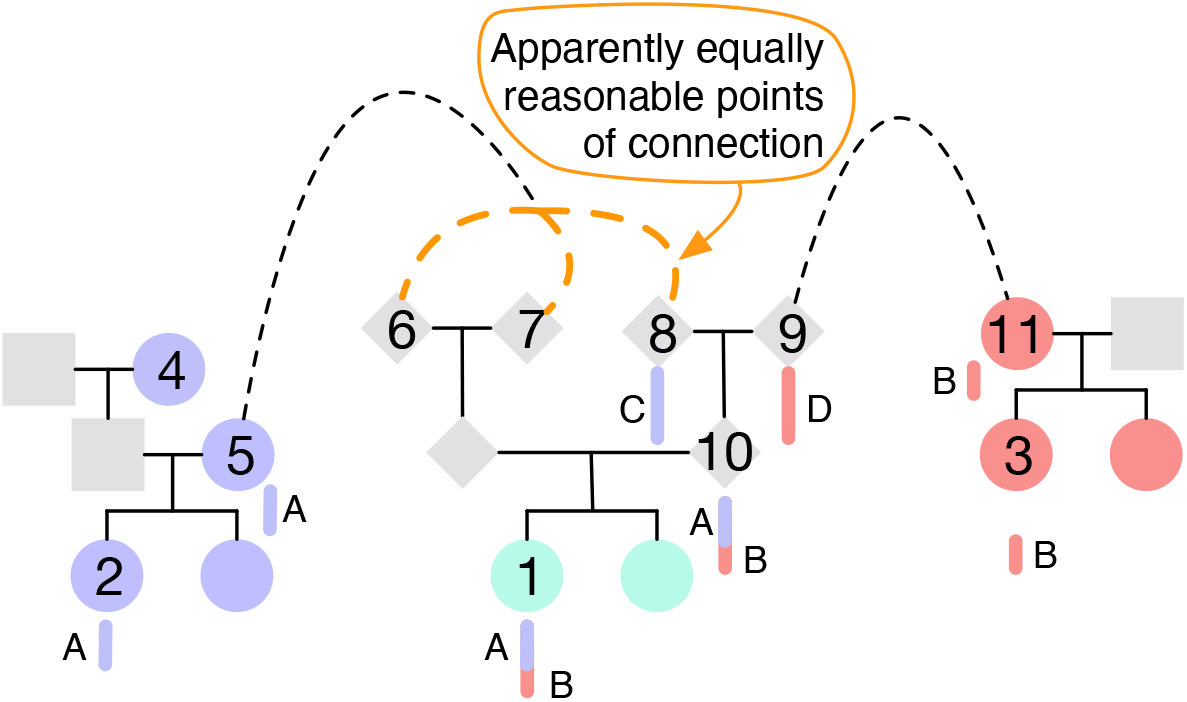
Determining the parental side of distant relatives. Individual 1 in the cyan pedigree shares segment *A* IBD with individuals 2 and 5 in the purple pedigree and they share segment *B* IBD with individuals 3 and 11 in the red pedigree. If the lineage connecting individual 1 to the purple pedigree passes through ancestor 8 and the lineage connecting individual 1 to the red pedigree passes through individual 9, then the ranges of segments *A* and *B* cannot overlap because individual 10 only transmits one recombined haplotype to individual 1. Observing abutting segments *A* and *B* is evidence that the cyan pedigree is connected to the purple and red pedigrees through the same parent. Observing spatially overlapping segments *A* and *B* is evidence that the purple and red pedigrees are connected through different parents of individual 1. In the absence of segment overlaps and splicing information, the orange dashed lines indicate equally reasonable ways to connect the purple and cyan pedigrees.

In the Big Bonsai method, when there are multiple possible grandparents of a common ancestor through which we can connect a focal set of nodes *𝒩* in a focal pedigree *𝒫* to two distantly-related pedigrees *𝒫*_1_ and *𝒫*_2_, we examine whether the IBD segments between *𝒫*_1_ and *𝒩* overlap with the IBD segments between *𝒫*_2_ and *𝒩*. The efficacy of checking segment overlaps is discussed in Section 4.3 using simulated data.

#### 3.5.7. Summary of the Big Bonsai algorithm

We combine the tools in Sections 3.5.2 – 3.5.6 to obtain the Big Bonsai method presented in Algorithm 4. The input for the Big Bonsai method consists of small pedigrees inferred using the Small Bonsai method. It assembles these small pedigrees into a large and sparsely-sampled pedigree by iteratively combining the two pedigrees that share the greatest total length of IBD until all pedigrees have been agglomerated into a single pedigree, or discarded because they cannot be combined in a reasonable way.

We assume that a pair of pedigrees, *𝒫*_1_ and *𝒫*_2_, can only be combined in ways that connect individuals who share IBD. When combining two pedigrees, the Big Bonsai method identifies the sets, *𝒩*_1_ and *𝒩*_2_ of genotyped nodes in each pedigree that share at least one IBD segment with an individual in the other pedigree. If the set *𝒩*_*i*_ does not have at least one common ancestor, we find the set *Ã*_*i*_ of most recent ancestral nodes whose descendants comprise *𝒩*_*i*_. The pair of ancestors *A*_1_ *∈Ã*_1_ and *A*_2_ *∈ Ã*_2_ whose descendants share the greatest total length of IBD is then determined and we redefine *𝒩*_1_ and *𝒩*_2_ to be the genotyped descendants of *A*_1_ and *A*_2_, respectively.

Our objective is to identify pairs of individuals through which *𝒫*_1_ and *𝒫*_2_ can be connected in such a way that all individuals in *𝒩*_1_ are related to all individuals in *𝒩*_2_. This is accomplished if and only if the sets *𝒩*_1_ and *𝒩*_2_ share at least one common ancestor. Sets *𝒩*_1_ and *𝒩*_2_ will be connected through a common ancestor if their respective common ancestors, *A*_1_ and *A*_2_, share a common ancestor or if *A*_1_ is descended from any individual in *𝒩*_2_ or from any ancestor on the induced subtree Λ_2_ of pedigree *𝒫*_2_ relating *𝒩*_2_ to one another. Similarly, sets *𝒩*_1_ and *𝒩*_2_ will have a common ancestor if *A*_2_ is descended from any individual in *𝒩*_1_ or from any ancestor on the induced tree Λ_1_ of pedigree *𝒫*_1_ relating *𝒩*_1_.

We present a generalized DRUID estimator in Appendix 6.3 for connecting pedigrees through individuals *A* who are not common ancestors of *𝒩*_1_ or *𝒩*_2_. However, connecting pedigrees *𝒫*_1_ and *𝒫*_2_ through all possible pairs can be computationally inefficient. Instead, we can accept a certain loss in accuracy and allow pedigrees to be connected only through common ancestors. We find that this approach works well in practice, generating pedigrees that are nearly as accurate as those constructed by connecting *𝒫*_1_ and *𝒫*_2_ in all possible ways.

Let *A*_1_ be a most recent common ancestor of *𝒩*_1_ and let *A*_2_ be a most recent common ancestor of *𝒩*_2_. For each pair of possible ancestors (*A*_1_, *A*_2_), we compute the generalized DRUID estimate *d*_*D*_(*A*_1_, *A*_2_) of the degree using Equation (9). We then perform the test for background IBD described in Section 3.5.5, which potentially results in a new pair of common ancestors 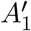 and 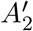 whose descendants do not share detectable background IBD. If the pair 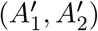 differs from the original pair (*A*_1_, *A*_2_), we replace *A*_1_ and *A*_2_ with 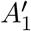 and 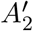 and recompute the generalized DRUID estimate *d*_*D*_(*A*_1_, *A*_2_). At the end of these steps, we have a set of possible ancestral pairs through which *𝒫*_1_ and *𝒫*_2_ can be connected, along with point estimates, *d*_*D*_(*A*_1_, *A*_2_), of the total degree separating each pair.

It remains to evaluate the likelihood of each pair and degree. Following the notation of Ko and Nielsen (2017), denote the relationship between a pair of individuals *A*_1_ and *A*_2_ with common ancestor (or ancestral pair) *G* by (*d*_1_, *d*_2_, *n*), where *d*_1_ is the number of meiotic events separating *A*_1_ from *G, d*_2_ is the number of meiotic events separating *A*_2_ from *G*, and *n* = |*G*| is the number of common ancestors. For a given estimate *d*_*D*_(*A*_1_, *A*_2_) of the degree between *A*_1_ and *A*_2_ and a number of common ancestors *n*, we consider all relationship types (*d*_1_, *d*_2_, *n*) corresponding to degree *d*_*D*_(*A*_1_, *A*_2_); in other words, we consider all relationship types such that *d*_1_ + *d*_2_ = *d*_*D*_(*A*_1_, *A*_2_) + *n−* 1.

For a given pair of ancestors *A*_1_ and *A*_2_, and for each relationship (*d*_1_, *d*_2_, *n*), we connect *A*_1_ to *A*_2_ through all such relationships and we evaluate the composite likelihoods of the resulting pedigrees computed using Equation (4). All pedigrees whose likelihoods are at least a fraction *f*_*𝓁*_ of that of the most likely pedigree are stored and the rest are discarded. We also apply the test in Section 3.5.6 for incompatible ancestral lineages to each retained pedigree and we retain only those pairs that pass the test.

Here, we have considered the procedure for combining two pedigrees *𝒫*_1_ and *𝒩*_2_. However, the output of the Small Bonsai method is a set of high-likelihood pedigrees *S* and the input to the Big Bonsai method is a list 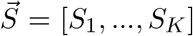 of such sets. Let *𝒩*_*S*_ denote the genotyped node set corresponding to the pedigree set *S*; in other words, *𝒩*_*S*_ is the genotyped node set of every pedigree *𝒫* ∈ *S*. If *𝒩* is the set of genotyped nodes in the full pedigree, then 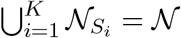.

At each step of the Big Bonsai method, we compare each pair of genotyped sets 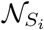 and 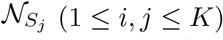 to determine the pair with the greatest shared total amount of IBD. Here, the total amount of IBD is the total length of IBD obtained by merging the segments shared between all pairs of individuals 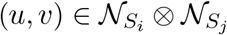. We then identify the subsets 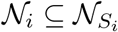 and 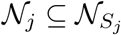 that share IBD and we combine each pair of pedigrees (*𝒫*_*i*_, *𝒫*_*j*_) ∈ *S*_*i*_ ⊗ *S*_*j*_ through all pairs of possible most recent common ancestors of *𝒩*_*i*_ and *𝒩*_*j*_. The full algorithm is presented in Algorithm 4.

It is possible to mis-infer relationships early in the process of pedigree building that lead to conflicts several steps later in the process. The downstream effects of a misplaced individual can be difficult to predict and prevent without a bird’s-eye view of the pedigree, but misplaced pairs of relatives can often be detected after the pedigree is built. In practice, we include a final step in the pedigree building process to detect internal inconsistencies by comparing the final pairwise relationships implied by the pedigree structure to the initial pairwise likelihood predictions. When the inferred relationships have low pairwise likelihoods, we rebuild the pedigree, iteratively expanding the number of pedigrees that are retained at each step to increase the chances that the correct pedigree is explored. We also correct pairwise point estimates that are likely to be incorrect when viewed in the context of a fully-built pedigree before attempting to re-infer the pedigree. Putting together the point estimator, the Small Bonsai method, and the Big Bonsai method, we obtain the full Bonsai method shown in Figure 1. Outlines of the three primary stages of Bonsai are shown in Algorithms 1, 2, and 4. The Bonsai method performs these stages in series.

### 3.6. Subjects and simulations

Our empirical analyses are based on simulated data, as well as a dataset comprised of the pedigrees of 23andMe research participants. All simulations and analyses that used real genotype data were performed using individuals consented for research at 23andMe.

#### 3.6.1. Overview of simulations

Simulations were carried out using two different general approaches. In one approach, no genotype or customer data were used and IBD segments were known with certainty, their positions and lengths being recorded during the simulation process. In the second simulation approach, the full-genome genotypes of research-consented 23andMe customers were used for the pedigree founders and genotypes were simulated for individuals in all subsequent generations through cross-over events. IBD segments were then inferred between each pair of individuals using

##### Algorithm 1 Pairwise likelihoods and point estimates.

Compute the likelihood of many different relationships between a pair of individuals, *i* and *j*, and obtain a point estimate of the relationship between *i* and *j*.

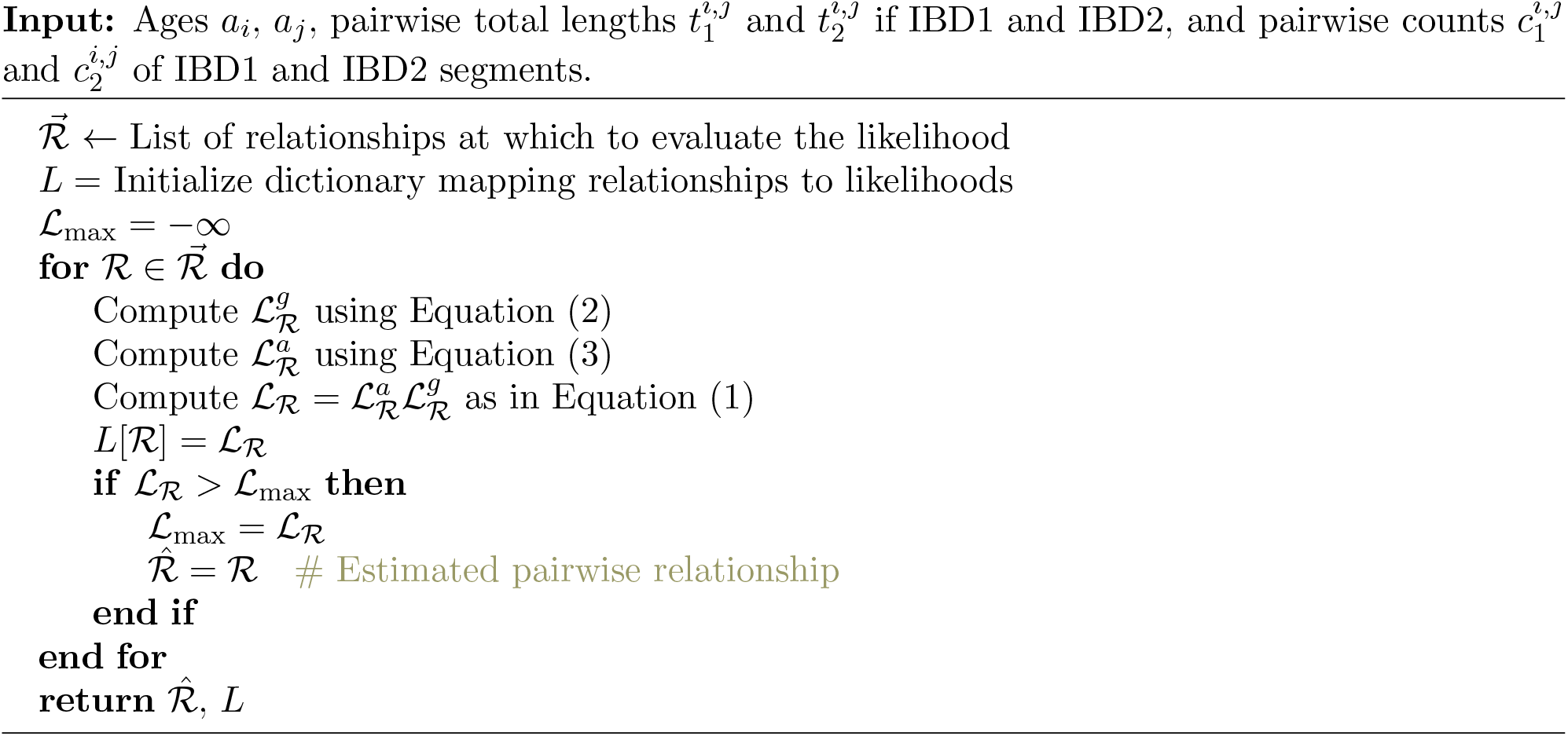

an in-house method for inferring IBD on unphased data (Henn et al. 2012), which is similar to that of Seidman et al. (2020)

In all simulations, the number of cross-over events in each meiosis was drawn such that the expected number of events was one per 100 cM and the locations of cross-overs were sampled uniformly along chromosomes.

#### 3.6.2. Validated real pedigrees

To evaluate Bonsai on true pedigrees, we constructed 204 validated pedigrees for individuals in the 23andMe database. By considering pedigrees in which all individuals were genotyped, we were able to construct each pedigree with a high degree of certainty by connecting parent-child pairs inferred using Algorithm 1. To ensure that the true pedigree was known with certainty, we considered quartets of genotyped customers with at least two full-sibling children and two parents. We identified pedigrees in which each individual was connected to every other individual through a chain composed of these building blocks. We further restricted our attention to pedigrees that spanned at least three generations with at least one pair of first cousins.

Pedigrees identified in this way allowed us to know the true pedigree structure because parent-offspring and full-sibling pairs can be inferred with nearly perfect accuracy and the quartet structure allows us to further confirm each inferred relationship using the other pairs in the quartet. In particular, we required that each sibling pair had inferred child-parent relationships with the same two parents using Algorithm 1. We also required the self-reported ages of both parents to be at least 17 years older than the self-reported ages of the children.

#### 3.6.3. Self-reported pedigrees

The Family Tree feature provided by 23andMe allows users to edit and validate relationships in their pedigrees. We considered a set of such pedigrees where users had either verified or changed relationships, indicating that they knew the correct relationships for at least a subset of individuals in the pedigree. We considered only individuals in these pedigrees who

##### Algorithm 2 Small Bonsai algorithm

Infer a Small Bonsai pedigree.

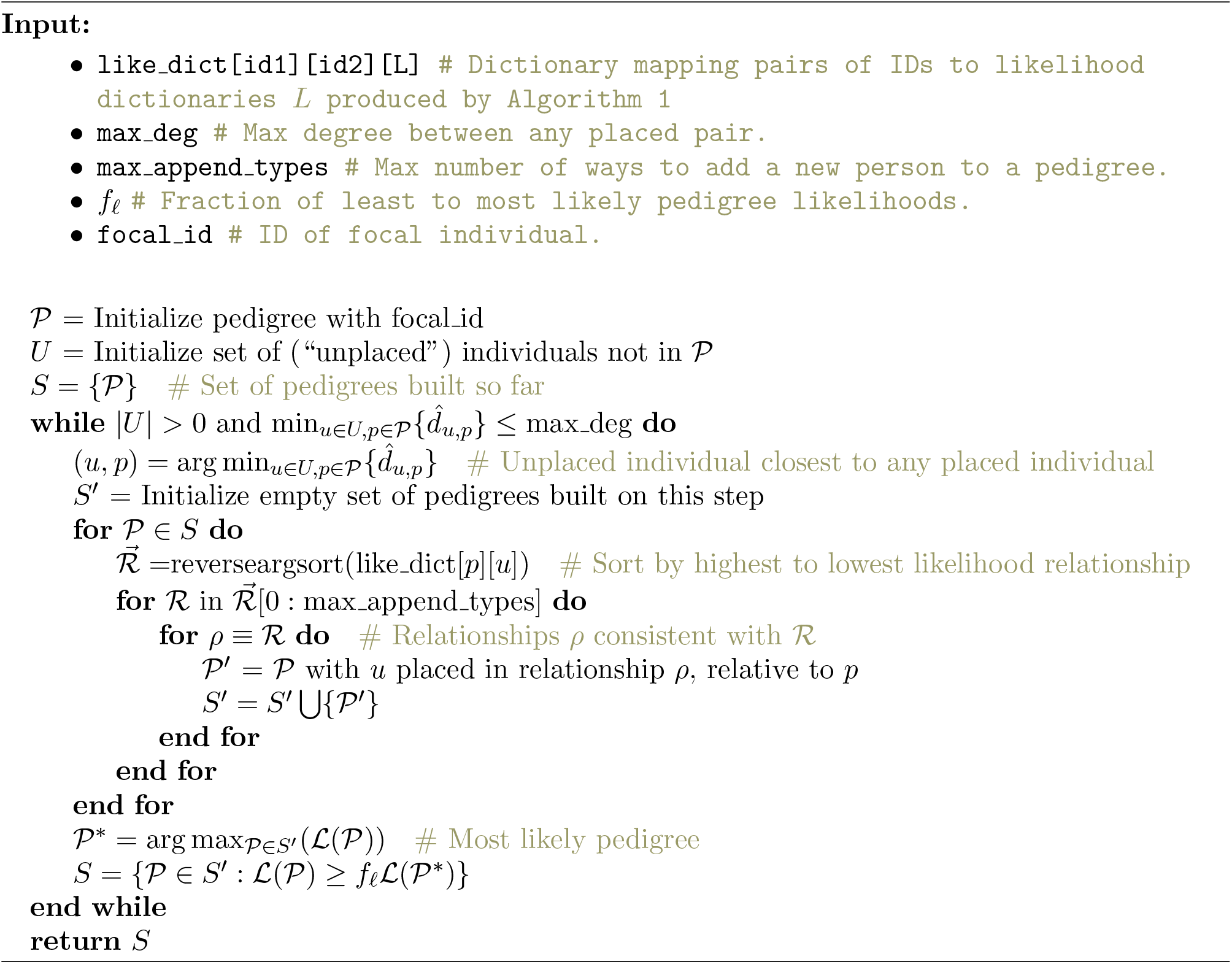

##### Algorithm 3 Detect background IBD

Detect whether the IBD observed in one of the clades directly descended from *A*_1_ in pedigree *𝒫*_1_ carries background IBD relative to the descendants of *A*_2_ in pedigree *𝒫*_2_.

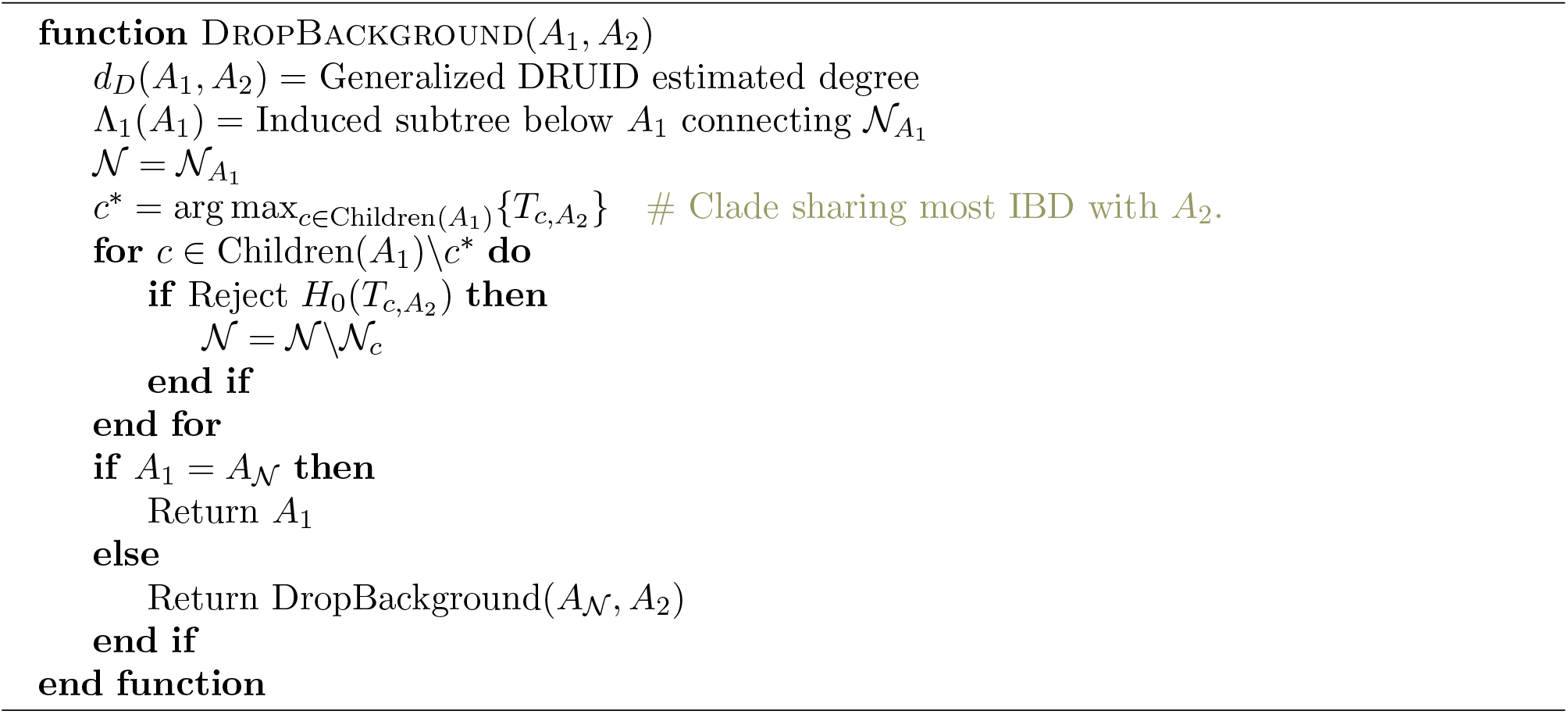

##### Algorithm 4 Big Bonsai algorithm

Combine Small Bonsai pedigrees into a Big Bonsai pedigree.

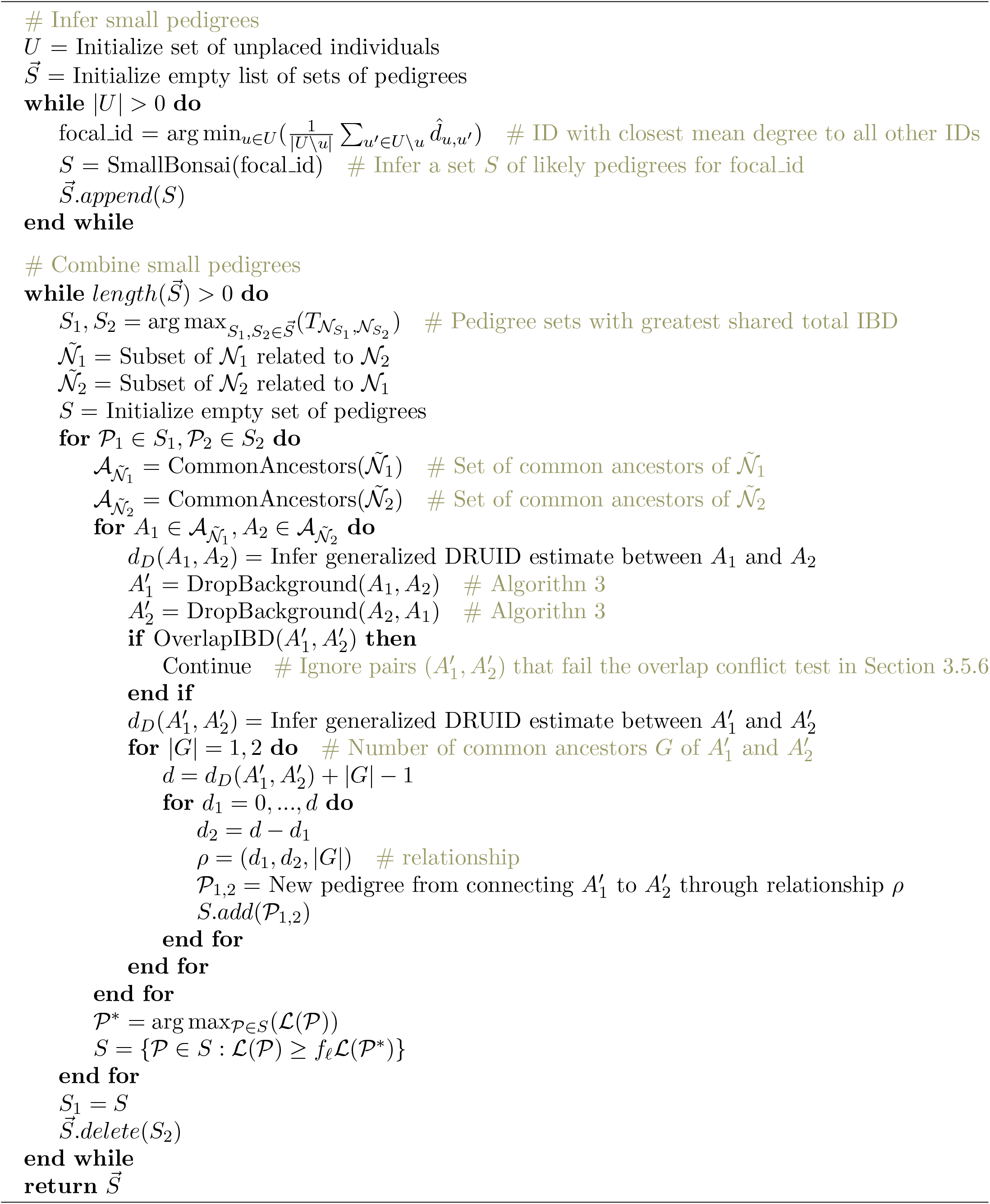

were consented for research and re-built the pedigree using only the subset of research-consented individuals. The inferred relationships in the pedigree could then be compared with the user-verified relationships.

#### 3.6.4. Simulations for fitting empirical pairwise genetic likelihood distributions

The distribution of the total length of IBD1 and IBD2, the distribution of lengths of IBD1 and IBD2 segments, and the distribution of the total counts of IBD1 and IBD2 segments for a specified relationship type *R* were obtained by simulating full genomes for 100 pairs of individuals of the relationship type. For each simulation replicate, a pedigree was specified containing the relationship of interest and cross-over events were simulated within the pedigree.

Over the 100 replicates, we computed the mean *µ*_*Q*_ and standard deviation *σ*_*Q*_ of the quantities *Q* = *T*_1_, *T*_2_, *C*_1_, and *C*_2_ where *T*_1_ is the total genome-wide length of IBD1, *T*_2_ is the total genome-wide length of IBD2, *C*_1_ is the total genome-wide count of IBD1 segments, and *C*_2_ is the total genome-wide count of IBD2 segments.

#### 3.6.5. Large simulated pedigrees

The 204 validated customer pedigrees described in Section 3.6.2 are small enough that the Small Bonsai method is capable of building them without resorting to the Big Bonsai method. To evaluate the Big Bonsai method, we required considerably larger pedigrees whose structures were known with certainty. Although many pedigrees for 23andMe research-consented customers are large, the relationships within them are typically not known with certainty. Therefore, we simulated large pedigrees to evaluate the Big Bonsai method.

Exact IBD was simulated for pedigrees with a depth of five generations by choosing a focal individual and building the “cone” of ancestors comprised of two parents, four grandparents, eight great-grandparents, and sixteen great-great-grandparents. For each individual in the ancestral cone, a second spouse was added with probability 0.2. Then, two children were created for every pair of spouses in the pedigree. Two children were repeatedly sampled for every spouse pair with no children until the generation with the focal individual was reached. An example of a pedigree generated by this approach is shown in Supplemental Figure S1.

#### 3.6.6. Simulated pedigrees for testing degree inference

The approach for simulating pedigrees for degree inference was similar to that in Section 3.6.5; however, the pedigree structure was different. For these pedigrees, we were interested in inferring the degree between a pair of common ancestors *A*_1_ and *A*_2_, given IBD observed between their descendants *𝒩*_1_ and *𝒩N*_2_.

For this analysis, we created two identical small pedigrees *𝒫*_1_ and *𝒫*_2_. Each small pedigree had the same structure comprised of the common ancestor *A*_1_ or *A*_2_, their spouse, their two children, and four grandchildren, where the grandchildren were comprised of two children for each child of *A*_1_ or *A*_2_. The ancestors *A*_1_ and *A*_2_ were then connected by degree *d*(*A*_1_, *A*_2_) through a pair of common ancestors, where the degree *d* varied from 1 to 10.

## 4. Results

We considered both simulated and real data to investigate the performance of the small and big Bonsai methods and their components.

### 4.1. Degree estimation

To evaluate the accuracy of degree inference using the likelihood estimator (Equation 16) and the generalized DRUID estimator (Equation 9), we applied these estimators to infer the degree between common ancestors *A*_1_ and *A*_2_ of two small pedigrees *𝒫*_1_ and *𝒫*_2_ (Section 3.6.6). Figure 8 shows the accuracy of the likelihood estimator *d*_*L*_ and the generalized DRUID estimator *d*_*D*_ for inferring the degree *d*, conditional on the event that any IBD at all was observed between the leaf nodes in *𝒫*_1_ and *𝒫*_2_. From Figure 8 it can be seen that both the maximum likelihood estimator *d*_*L*_ and the generalized DRUID estimator *d*_*D*_ have similar accuracies for inferring the degree *d*. Moreover, the DRUID estimate is nearly identical to the maximum likelihood estimate, which is important in practice because it implies that connecting two pedigrees through the degree inferred by DRUID results in a pedigree that is approximately the maximum likelihood pedigree. This result can dramatically speed up pedigree inference and, in practice, we use the generalized DRUID estimator for inferring the degree of separation between two small pedigrees.

**Figure 8.**
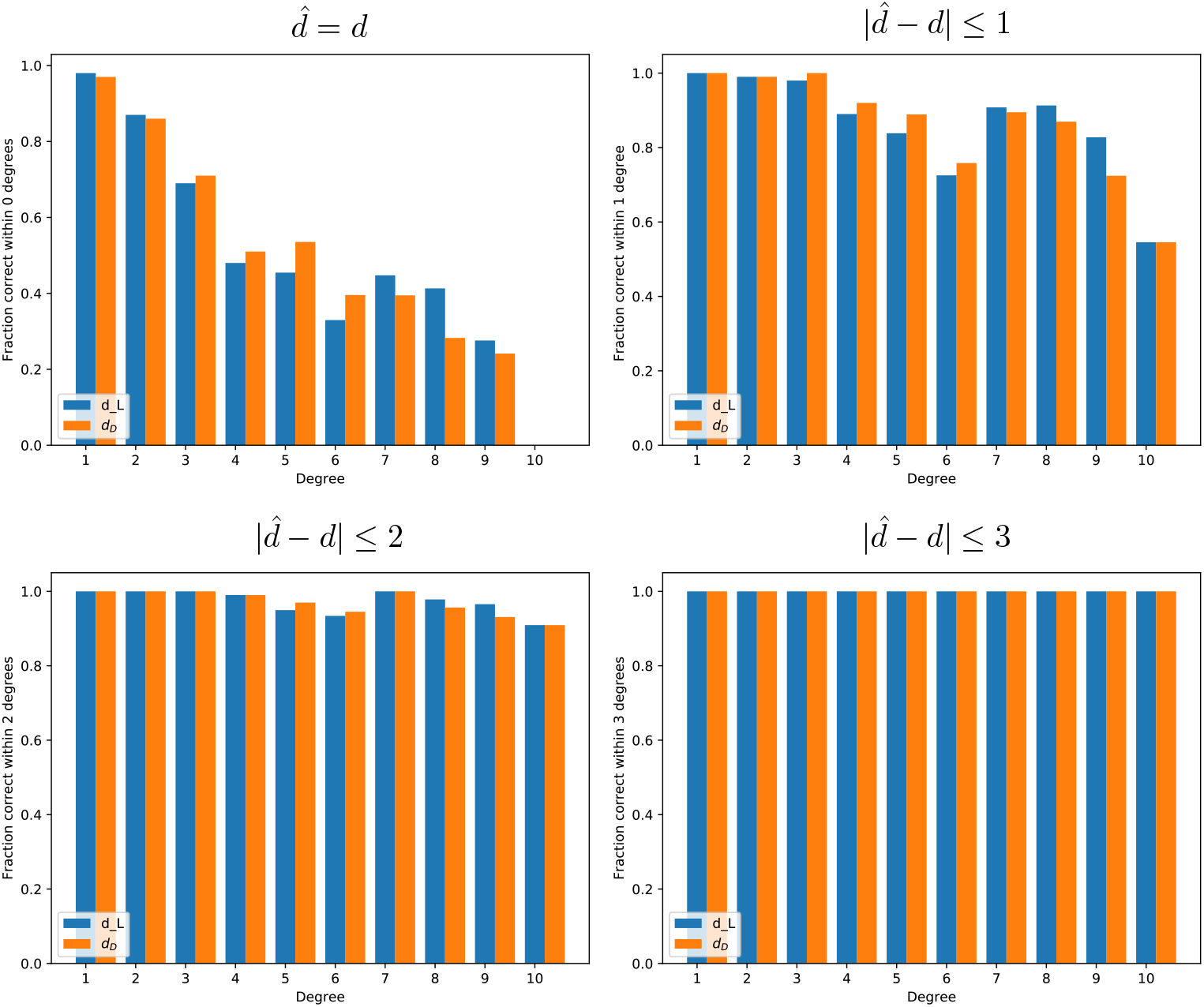
The accuracy of the likelihood method (Equation 16) and the generalized DRUID method (Equation 9) for inferring the degree between a pair of common ancestors. The accuracy of the estimate is shown for four different tolerances: exactly equal to the true degree, within one degree of the true degree, within two degrees of the true degree, and within three degrees of the true degree.

### 4.2. Background IBD detection

To evaluate the efficacy of the test in Equation (17) for detecting background IBD, we simulated pedigrees comprised of three small pedigrees, *𝒫*_1_, *𝒫*_2_, and *𝒫*_3_, connected together (Figure 9). In one set of simulations, pedigree *𝒫*_1_ was related only to *𝒫*_2_ and not *𝒫*_3_ (Figure 9, Scenario 1). This allowed us to simulate background IBD among all pairs of individuals and then attempt to detect it. In another set of simulations, pedigree *𝒫*_1_ was truly related to all other individuals (Figure 9, Scenario 2). This second set of simulations allowed us to evaluate the rate at which background IBD was detected even when there was true IBD between *𝒫*_1_ and *𝒫*_3_ as well as background IBD. Note that in all simulations, all pairs shared a nonzero expected amount of background IBD so that even truly related individuals carried additional background IBD.

**Figure 9.**
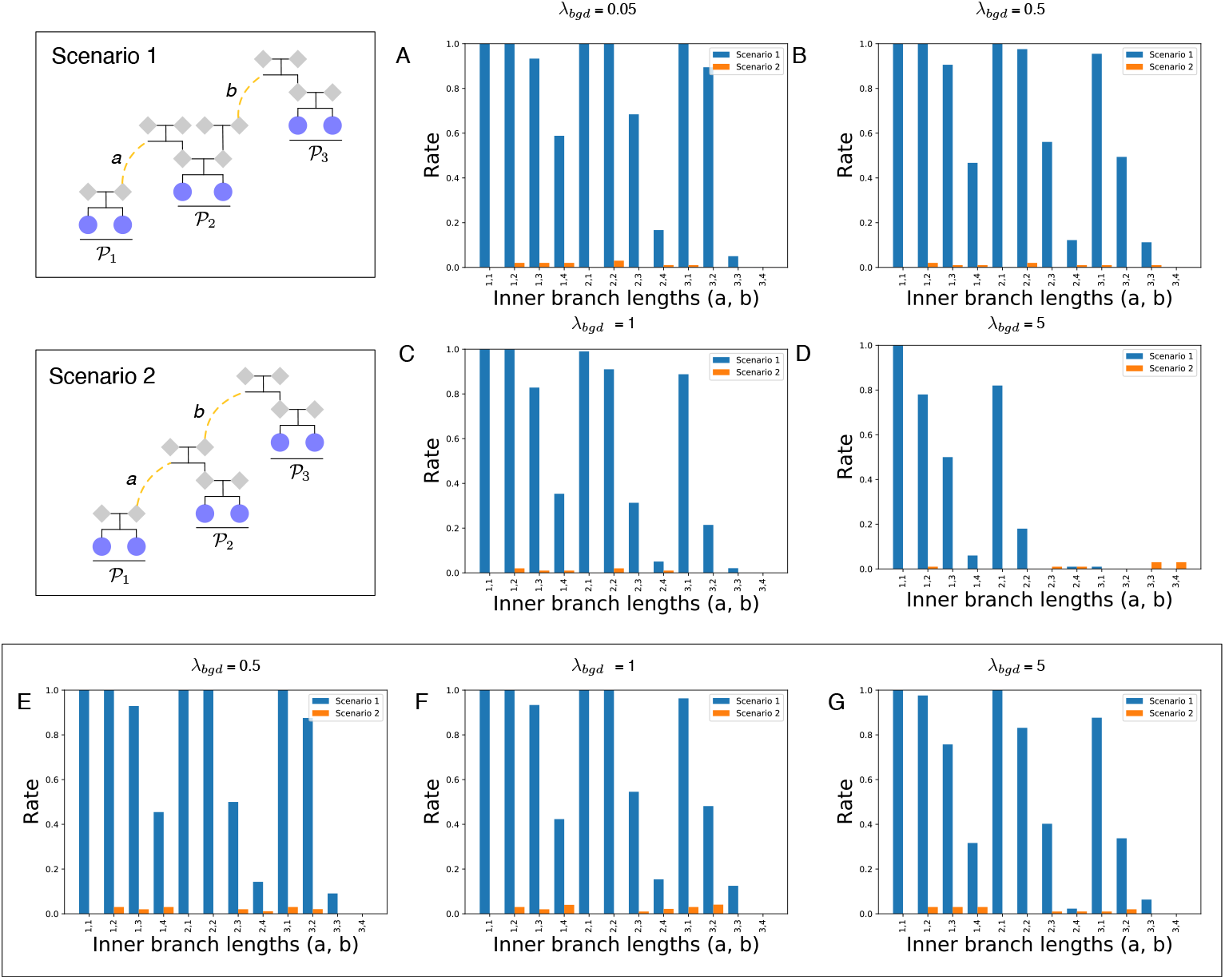
Evaluating the use of the test in Equation (17) for detecting background IBD. Scenario 1 shows a pedigree structure in which any IBD observed between *𝒫*_1_ and *𝒫*_3_ is background IBD. Scenario 2 shows a pedigree structure in which IBD shared between *𝒫*_1_ and *𝒫*_3_ comprises both true and background IBD. Branch lengths *a* and *b* were variable. (A)-(D) Rates for detecting background IBD under scenarios 1 and 2. (E)-(G) Same as (B)-(D), but using only segments at least 10 cM in length. Plots are shown for *α* = 10^−3^.

For all pedigrees, we simulated background IBD between each pair of individuals by sampling the number of background IBD segments from a Poisson distribution with mean *λ*_*bgd*_ = 0.05, 0.5, 1, or 5. We then sampled the length *L* of each observed background segment from a thresholded exponential distribution with mean 7 cM and minimum length of 5 cM. A minimum of 5 cM was chosen because, in practice, small segments can be difficult to infer and it is a common practice to employ a minimum cutoff on the length of IBD segments to reduce the rate of false positives (Huff et al. 2011).

The rates of background IBD we tested corresponded to values spanning the empirically observed range of background segment counts in broad human populations in the 23andMe database. When considering only segments greater than 5 cM in length, the average number of background IBD segments between a pair of individuals is between 0.01 and 0.02 for most human populations. However, for populations with historical bottlenecks, the expected background IBD count can be closer to *λ*_*bgd*_ = 5.

Figure 9A shows the fraction of times the null hypothesis *H*_0_ in Equation (17) was rejected when individuals shared an average of 0.05 background IBD segments. Blue bars correspond to simulation replicates in which all IBD shared between *𝒫*_1_ and *𝒫*_3_ was background (Scenario 1) and orange bars correspond to simulation replicates in which background IBD between *𝒫*_1_ and *𝒫*_3_ existed in addition to true IBD.

Each bar in Figure 9A was calculated using 50 simulated pedigrees. From Figures 9A and 9B, it can be seen that for a level of background IBD consistent with the majority of human populations, the test correctly identified background IBD a large fraction of the time. Moreover, the test typically did not detect background IBD when there was also true IBD in addition to background IBD.

For high levels of background IBD consistent with populations with severe bottlenecks, it was much more difficult to detect background IBD (Figures 9C and 9D). This was especially true when the internal branch length *b* was long and background IBD dominated true IBD.

This is problematic because the detection of background IBD is particularly important in these populations. However, the amount of background IBD is controllable to some degree by establishing a threshold for the minimum length of an IBD segment to be included in an analysis. The higher the threshold, the fewer the number of false positive segments and the longer the length of the background IBD segments that are not filtered by the threshold. By using this threshold when detecting background IBD, it is possible to increase the power for detecting background IBD. This can be seen in Figures 9E-G, which show the same simulations shown in 9B-D, but discarding all segments shorter than 10 cM.

### 4.3. Segment overlap detection

We evaluated the degree to which overlapping IBD segments can be informative about the ancestors through which two pedigrees are connected using the large simulated pedigrees described in Section 3.6.5. For each pedigree, we considered the four grandparents of a focal individual and the leaves descended from all lineages extending up from each of the four grandparents. In the example large pedigree shown in Figure S1, the focal individual is one of the yellow leaf nodes and the clades corresponding to the four leaf sets are colored in green, cyan, red, and magenta.

For a pair of leaf sets related to the focal individual through an ancestral couple, we expect to see no overlap in the IBD segments shared with the focal individual. For a pair of leaf sets related to the focal individual through two grandparents who are not a couple, we expect to observe overlapping segments occasionally.

Figure 3 shows the rate at which segments from one leaf set overlapped segments from another leaf set by more than a fraction *f* of the total IBD observed between the two leaf sets, combined, for *f* ∈ 0.01, 0.05, 0.1, 0.15, 0.2.

Let *i* denote the focal individual. For leaf sets *𝒩*_1_ and *𝒩*_2_ with total amounts of IBD to the focal individual denoted by 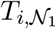 and 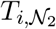, let 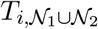 denote the total length of merged segments between focal individual *i* and either set. We recorded an overlap in segments if the following relationship was satisfied: 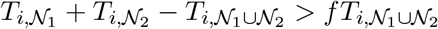.

Figure 10 indicates that even with few sampled leaves from each leaf set, it is possible to detect overlapping IBD segments a large fraction of the time when the leaves are related through grandparents who are not a couple. Each bar in Figure 3 was computed using 100 pedigrees, each with four pairs of leaf sets related to individual 1 through a pair of grandparents who were not a couple. Only IBD segments greater than 5 cM in length were considered.

**Figure 10.**
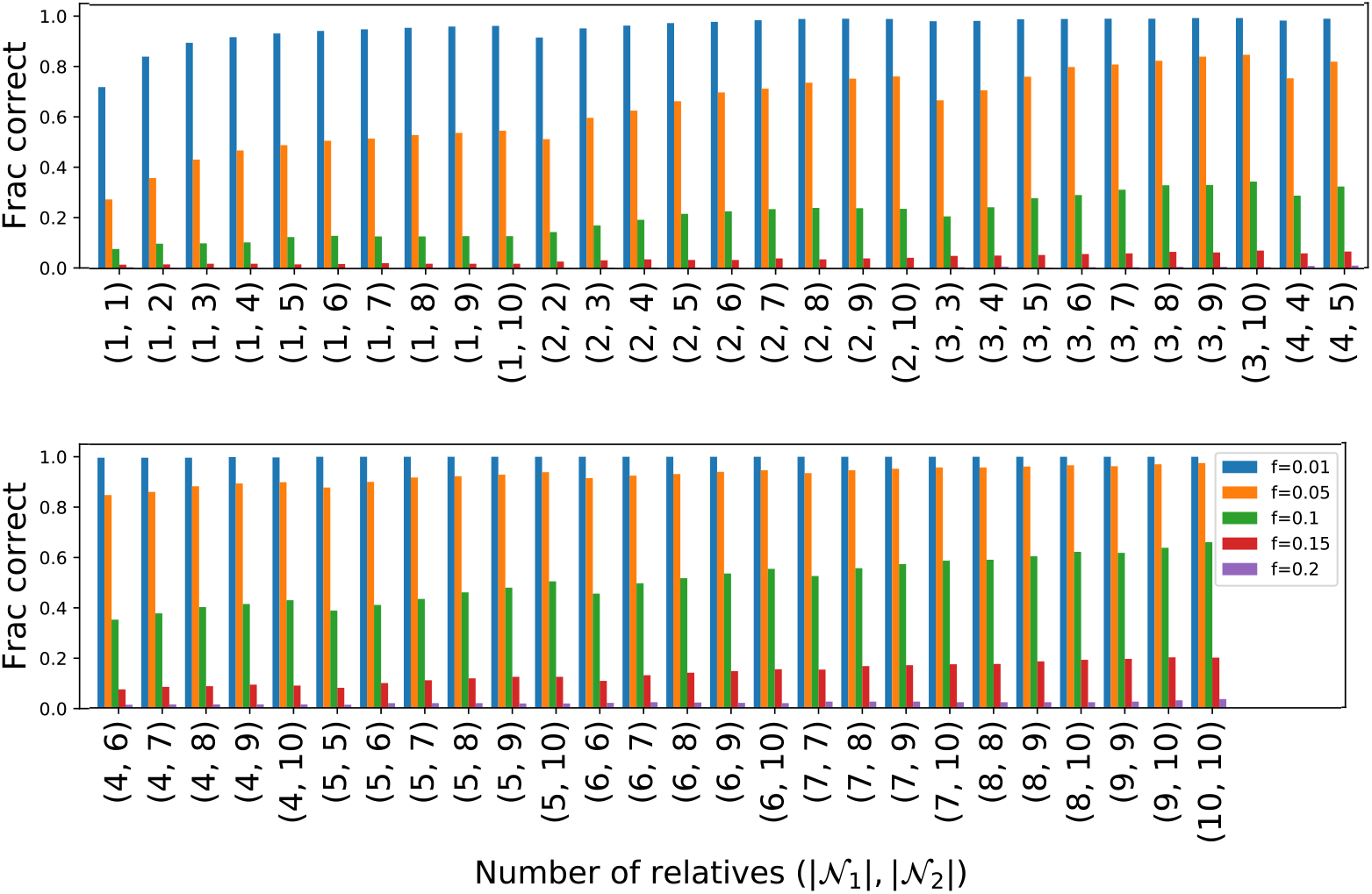
The probability of observing an IBD segment overlap. The plot shows the probability of observing an overlap of at least fraction *f* (*f* = 0.01, 0.05, 0.1, 0.2) among segments shared IBD between the focal individual and sets of leaves related to the focal individual through ancestors who are not a couple. IBD segments were simulated for large pedigrees like that shown in Figure S1. IBD was computed between the focal individual and the leaf nodes of each of the four clades related to the focal individual through each of the four grandparents (colored green, cyan, red, and magenta in Figure S1). An observed IBD segment overlap was evidence that the lineages were related to the focal individual through a pair of ancestors who were not a couple.

### 4.4. Timing and accuracy of Small Bonsai, compared with PRIMUS

To evaluate the accuracy and runtime of Bonsai in comparison with PRIMUS, we applied PRIMUS and Bonsai to a set of 204 pedigrees comprised of research-consented 23andMe customers (Section 3.6.2) in which all individuals were genotyped and for which the true pedigree was known with a high degree of certainty.

Pedigrees in which all individuals have been genotyped are simple to infer by connecting together first degree relatives. The difficulty is in constructing pedigrees in which only a small fraction of individuals have been genotyped. Therefore, to evaluate the accuracy of Bonsai, we subsampled the validated pedigrees and performed inference using the subset of individuals, ignoring the remaining individuals. The resulting pedigree could then be compared to the subgraph of the true pedigree corresponding to the subsampled individuals to determine the accuracy of the inference.

We subsampled each pedigree to 10, 20, 30, 40, 50, 60, 70, 80, 90, or 100% of its members with a minimum of at least two individuals sampled per pedigree. Figure 11 shows the degree to which each method recovered each relationship type. From Figure 11, it can be seen that the Bonsai algorithm achieved improved accuracy for inferring relationships, and that when at least 50% of individuals were sampled, Bonsai inferred the correct pedigree with near perfect accuracy.

**Figure 11.**
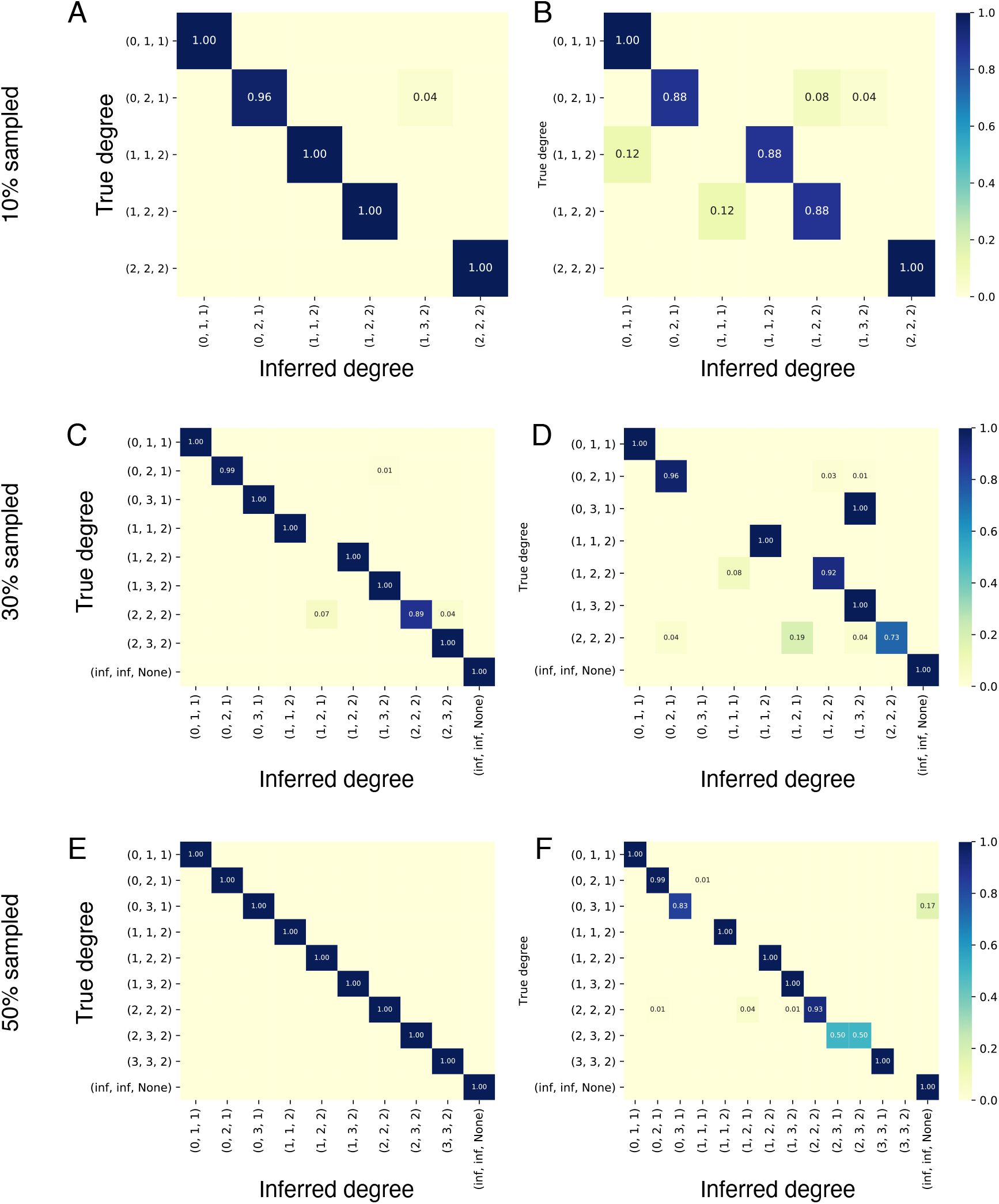
Comparison of Bonsai with PRIMUS. Panel A shows the accuracy of Bonsai for inferring different relationships when 10% of individuals were sampled from each pedigree. The relationship type between a pair of individuals *i* and *j* is indicated as a tuple of the form (*d*_*i,G*_, *d*_*j,G*_, |*G*|) following the notation of Ko and Nielsen (2017). Panel B shows the accuracy for PRIMUS applied to the same individuals with the same pairwise likelihoods as Panel A. Panels C and D compare Bonsai with PRIMUS when 30% of individuals were sampled and panels E and F compare Bonsai with PRIMUS when 50% of individuals were sampled. Accuracy of Bonsai was perfect for 60-100% of lineages, although PRIMUS continued to mis-infer some relationships.

We also compared the runtime of the Bonsai method to the runtime of PRIMUS for the same set of pedigrees described in Section 3.6.2. Figure 12 shows the runtime for Small Bonsai compared to the runtime for PRIMUS for different percentages of sampled lineages from each of the pedigrees. The runtimes for Bonsai and PRIMUS were similar when few or many individuals were sampled, although Bonsai was slightly faster. In these regimes, pedigrees were fast to construct because individuals could be connected through close relatives, which were inferred with high confidence. However, when the number of individuals sampled was moderate and there were several possible pedigree configurations with high likelihoods, the Bonsai method was significantly faster than PRIMUS.

**Figure 12.**
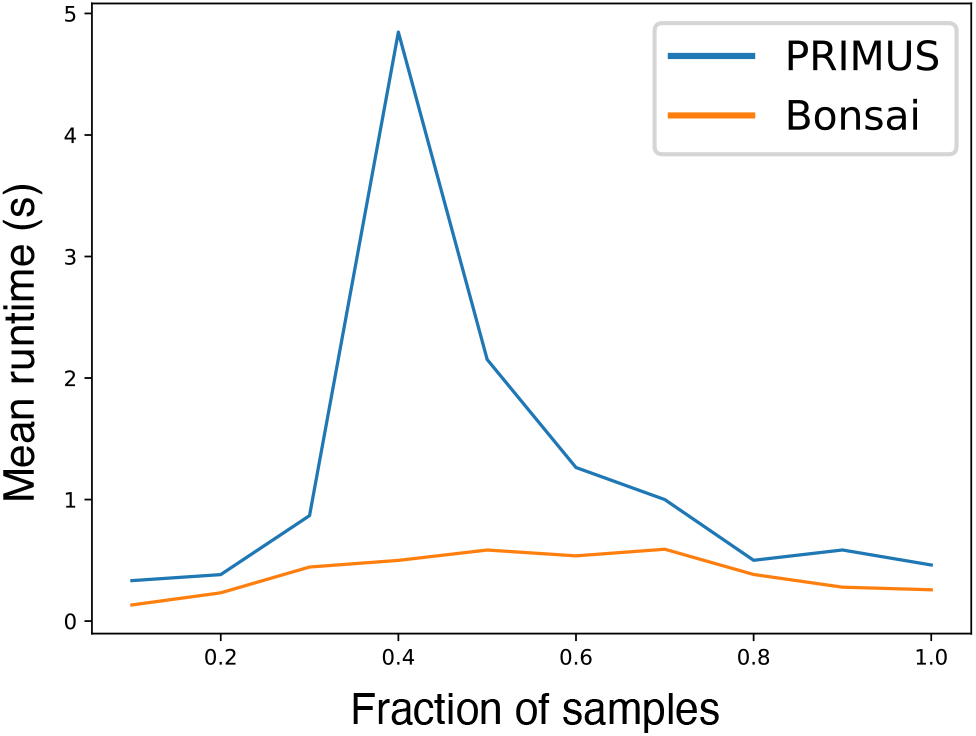
Comparison of runtime between Bonsai and PRIMUS. Runtime was evaluated using 204 pedigrees of 23andMe research participants that were known with a high degree of certainty. Because every individual in each pedigree was genotyped, we sampled 10, 20, 30, 40, 50, 60, 70, 80, 90, or 100% of their members uniformly at random without replacement. The subsampled individuals were then used to reconstruct the pedigrees using PRIMUS and Bonsai using the same IBD, age, and sex data. The x-axis in the figure is the fraction of sampled individuals in each of the 204 pedigrees that were used for pedigree inference. The y-axis is the average time in seconds required to reconstruct a pedigree.

### 4.5. Timing and accuracy of the Big Bonsai method

Reconstruction of large pedigrees using the Small Bonsai method can be computationally challenging due to a quickly-expanding state space of possible pedigrees. Figure 13 shows timing and accuracy results for reconstructing large five-generation pedigrees simulated using the approach described in Section 3.6.5. For these analyses, we were interested in the ability of Big Bonsai to infer pedigrees that were realistic representations of direct-to-consumer genetic data. In realistic pedigrees, individuals beyond the most recent two generations may not have been sampled. When sampling individuals for the pedigree, we sampled individuals only in the most recent two generations who were not founders. This provided a pool of approximately 130 individuals who could be sampled out of a total of at least 261 individuals in each pedigree, including pedigree founders.

**Figure 13.**
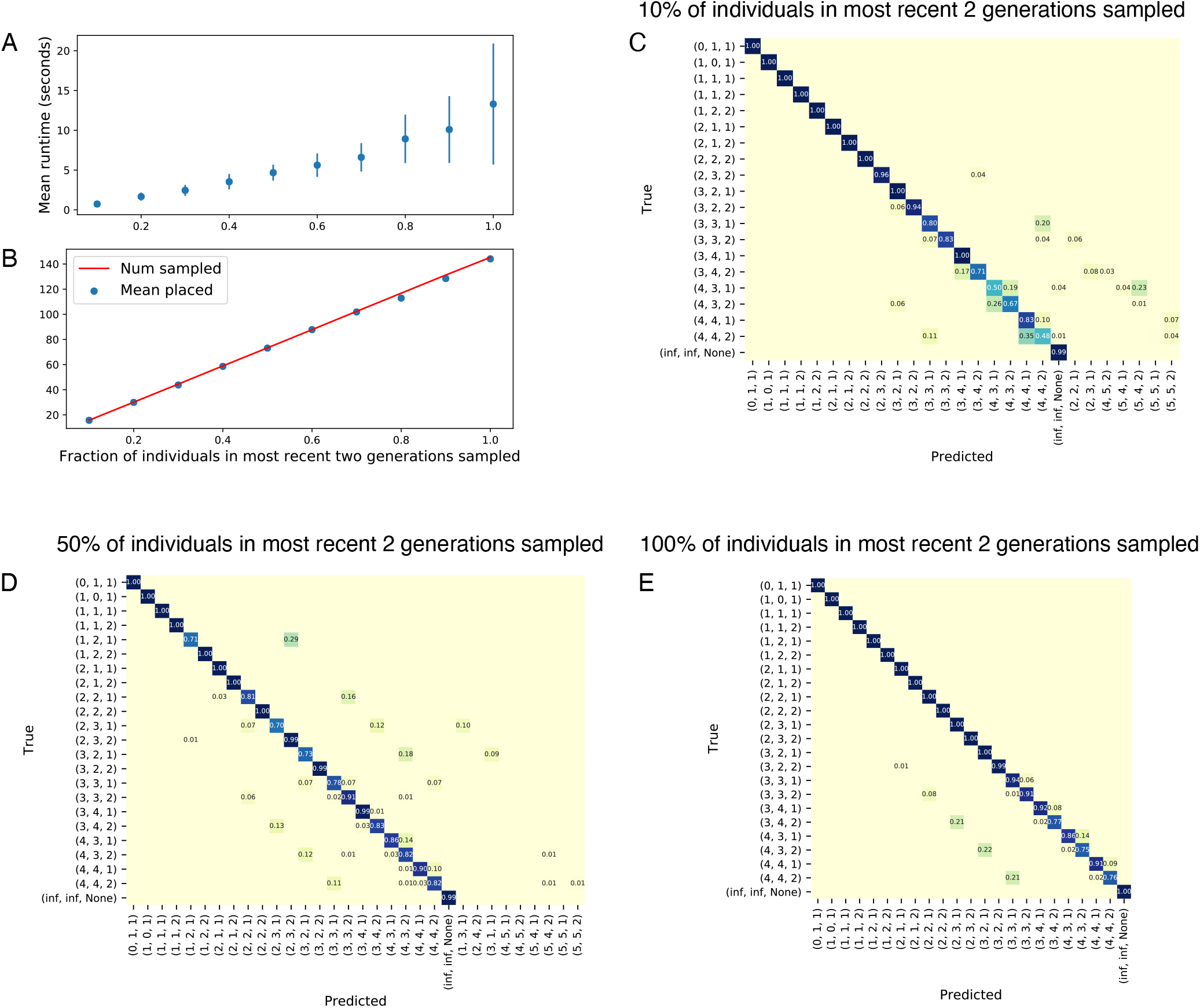
Timing and accuracy of the Big Bonsai method. Large pedigrees were simulated with a depth of five generations and two offspring per pair as described in Section 3.6.5. To capture the sparsity of pedigrees observed in direct-to- consumer customer data, we sampled only a fraction of individuals in each pedigree and used these as the genotyped individuals to infer the pedigree. Individuals were only sampled from the most recent two generations because ancestors are often ungenotyped in human data and because inference is more challenging when genotyped ancestors are unavailable. (A) Runtime for Big Bonsai as a function of the fraction of sampled individuals in the most recent two generations. (B) The number of sampled individuals in each pedigree and the mean number placed, averaged across 100 replicates. (C)-(E) The fraction of pairs with a given relationship type that were inferred to have each other relationship type. Tuples (*d*_*i,G*_, *d*_*j,G*_, |*G*|) indicate a specific relationship type between individuals *i* and *j* using the notation of Ko and Nielsen (2017): (up, down, number of common ancestors). The tuple (inf,inf,None) indicates unrelated individuals.

To evaluate the ability of the Big Bonsai method to reconstruct pedigrees with sparsely sampled individuals, we further subsampled 10, 20, 30, 40, 50, 60, 70, 80, 90, or 100% of the approximately 130 non-founder individuals in the most recent two generations. Sampling 10% of these individuals corresponds to sampling approximately 5% of all individuals in the full pedigree and sampling 100% of these individuals corresponds to sampling approximately 50% of all individuals in the pedigree overall. Our sampling scheme presents a further challenge to pedigree reconstruction because the samples did not contain ancestral individuals who could provide additional information about the degrees of distant relationships.

From Figure 13A, it can be seen that the runtime is on the order of several seconds per pedigree, even though pedigree sizes were large. Bonsai built pedigrees with over one hundred sampled individuals in tens of seconds.

The Big Bonsai method is designed to drop small pedigrees from consideration, rather than combining them with the other pedigrees when an inconsistency is detected. This can occur, for example, if the small pedigree is inferred with a very unlikely relationship despite re-running with parameter values that search a broader pedigree space and attempting to correct relationships that are judged to be inaccurate. Figure 13B indicates that the fraction of times individuals or small pedigrees were dropped was small, as the number of placed individuals was typically very close to the number of sampled individuals.

Figures 13C-D show the accuracy for inferring large pedigrees when different fractions of individuals were sampled. Close relationships were typically reconstructed accurately, whereas distant relationships were more challenging, yet still generally accurate especially when the fraction of sampled individuals was high.

Note that, because the ages of individuals in the pedigree conformed to average age differences between generations, it was sometimes possible to distinguish distant half relationships from distant full relationships. For example, a pair of individuals of the same age related by four degrees of separation is more likely to be a pair of half first cousins, rather than a full first cousin once removed. Half relationships are likely to be more challenging to infer in practice, given that age differences may differ from expectation.

### 4.6. Reconstruction of self-reported pedigrees using Big Bonsai

We also compared relationships inferred by Bonsai with self-reported relationships using 265 pedigrees for which the relationships between two or more individuals had been self-reported by the focal individual for whom the pedigree was built (Section 3.6.3).

Figure 14 shows the correspondence of each inferred relationship type with the self-reported relationship type. The plots show the fraction of times the self-reported and inferred relationships agreed exactly in that their relationship tuples (up, down, number of ancestors) were the same. The plots also show the fraction of times the relationships agreed in degree, the fraction of times the relationships agreed within one degree, and the fraction of times relationships agreed within two degrees.

**Figure 14.**
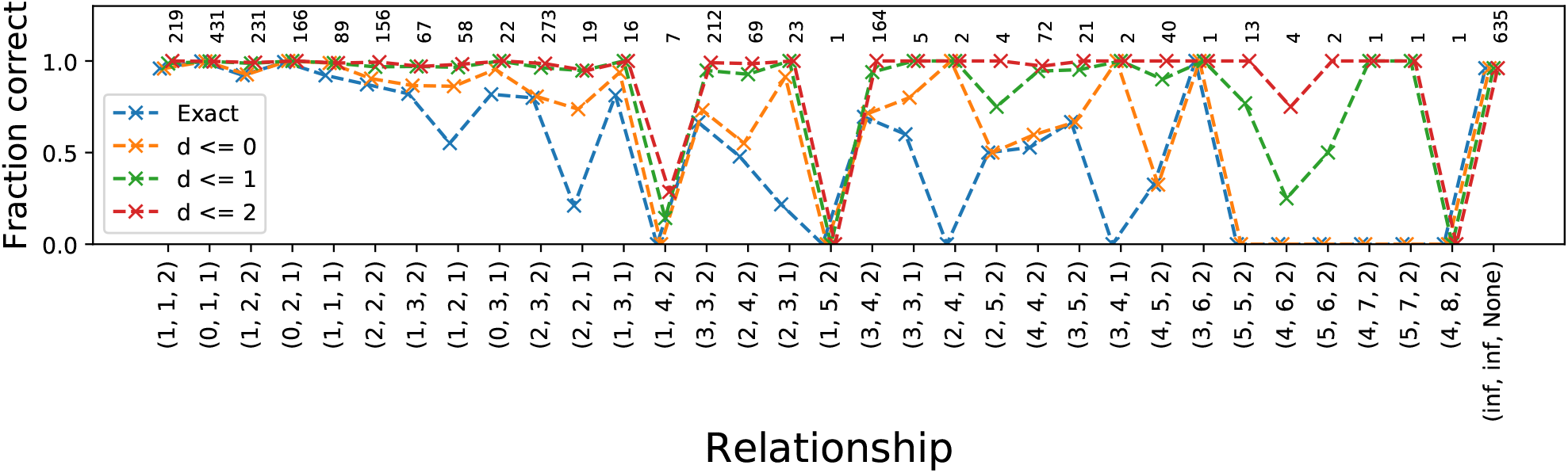
Comparison with self-reported pedigrees. Comparison of predicted relationships with self-reported relationships. Blue markers show the fraction of relationship pairs for which the inferred and self-reported relationships agreed exactly. The orange, green, and red markers show the fraction of pairs for which the degrees of the inferred and self-reported relationships differed by at most 0, 1, or 2 degrees, respectively. The number of pairs for each relationship is shown above the curves. Dashed lines are included to improve visibility.

The inferred and self-reported relationships typically agreed for close relationships up to first cousins. However, the inferred relationship often differed from the self-reported relationship for distant relationship types, and occasionally for relatives as close as siblings or parents. For parent-child and full sibling pairs, it is possible to check whether the self-reported relationship is correct because the IBD sharing patterns for these relationships are very distinct from other relationship types. It is of interest to note that in all but one case in which the inferred and self-reported relationships differed for a parent-child or full sibling pair, the self-reported relationship was, in fact, incorrect due to impossible levels of shared IBD. In these cases, it was frequently the case that a self-reported parent-child pair shared no IBD, or that a self-reported full sibling pair shared no IBD2 and instead had an IBD sharing pattern that was more consistent with a half sibling or a cousin. In only one case was the self-reported relationship type consistent with the IBD sharing pattern, and in this case one individual had a self-reported age much greater than 100 years, leading to a strong contribution from the age component of the likelihood and an incorrectly inferred relationship type.

For distant relationships, we observed greater disparities between the self-reported and inferred values. However, the inferred degree was often within one or two degrees of the self-reported relationship, even for relationships as distant as seventh degree or higher in some cases. Moreover, relationships for which the self-reported and inferred degrees differed by more than two degrees typically had few self-reported pairs (Figure 14). This relatively high accuracy for distant relationship degree is consistent with our analysis of the accuracy of the generalized DRUID estimator.

## 5. Discussion

We have presented a method for inferring large pedigrees quickly and accurately, even when the fraction of genotyped individuals in a pedigree is low and the distance between an individual and their closest relative can be moderate or large. Our method has three component algorithms that are applied in sequence: 1) a method to infer the likelihoods of pairwise relationships between each pair of individuals using both age and IBD data, 2) a method for inferring pedigrees of small-to-moderate size, and 3) a novel method for combining small pedigrees together into large and sparsely-sampled pedigrees.

Our Small Bonsai algorithm efficiently explores the space of possible pedigrees using a constructive approach. This approach is similar to that of PRIMUS (Staples et al. 2014), but it employs several new features that make it more efficient and more accurate than PRIMUS, including incorporating ages directly into the likelihoods, expanding the set of pedigrees that are explored, and introducing a branch-and-bound-like method for exploring the space of pedigrees more efficiently.

Although the new methodological approaches implemented in the Small Bonsai method provide a pedigree inference algorithm with improved accuracy and performance, the primary novelty of the Bonsai method is in the Big Bonsai algorithm, which combines small pedigrees together into large and sparsely-sampled pedigrees. This algorithm makes it possible to construct pedigrees that are much bigger than the maximum size that can be constructed by current approaches.

The construction of large and sparse pedigrees requires a fundamentally different approach from combining individuals one at a time as is done in PRIMUS, or searching a broad pedigree space by rearranging pedigrees as is done in CLAPPER. Because the space of possible pedigrees is large, it is useful to proactively and dramatically narrow the set of possible pedigrees to include only the pedigrees with the highest likelihoods.

Combining small pedigrees together into large and sparse pedigrees makes it possible to leverage information in the previously-inferred small pedigrees to identify the most likely ways in which the small pedigrees can be connected together. Leveraging information across small pedigrees allows us to more accurately infer the degree of relatedness between two small pedigrees and to identify background IBD.

We have introduced three new tools for combining pedigrees together. First, we have generalized the DRUID method of Ramstetter et al. (2018) to apply to general outbred pedigrees, rather than specific pedigree structures. We have also extended the method to allow pedigrees to be connected through pairs of individuals who are not common ancestors. We have also shown that the generalized DRUID estimate is nearly identical to the maximum likelihood estimate. Thus, rather than exploring multiple ways of connecting two pedigrees and selecting the most likely pedigree, we can simply connect the two pedigrees through the DRUID point estimate and achieve nearly the same result, greatly speeding up the inference process.

The second tool we have introduced is an approximate likelihood for the degree separating the common ancestors of two pedigrees as a function of the total length of IBD shared by the pedigrees. This likelihood is used as the foundation for our method for testing whether the IBD shared between two sets of individuals is the result of a true relationship, or whether the IBD is background IBD. Our approach obviates the need to infer the population or family-level distribution of IBD, which is useful because the expected amount of background IBD between a pair of individuals can be challenging to know in advance. By testing IBD between groups of individuals rather than pairs, we also reduce problems with multiple testing.

Although we intentionally did not incorporate the population or family-level distribution of background IBD into our background IBD detection method, one can imagine a method that combines our approach with such a distribution to improve the power for detecting background IBD when the population or family-level distribution of background IBD is known.

Finally, we have introduced a method for determining when the connection of pedigrees through certain ancestral branches is inconsistent with patterns of IBD overlap. This method makes it possible to assign two pedigrees to the correct parental sides of a focal individual in a focal pedigree. Using only information contained in pairwise IBD sharing, these inconsistent pedigrees would not be detected; pedigrees formed by connecting two pedigrees through incompatible grandparental lineages would appear to have the same likelihood as the true pedigree. Our approach achieves high accuracy even when few relatives on each parental side have been sampled.

Compared to previous methods for inferring complex human pedigrees, the Bonsai method yields improvements in both accuracy and computational efficiency and makes it possible to build pedigrees that are considerably larger than those that were possible before. The speed of pedigree building depends on the complexity of the pedigree, the proportion of individuals who are genotyped, and the distribution of these individuals throughout the generations of the pedigree. As a result, it can be difficult to characterize the runtime of Bonsai relative to other methods. However, in a comparison of runtime on 204 real-world pedigrees, Bonsai was always faster than the current fastest method PRIMUS. For large complicated pedigrees, Bonsai built pedigrees in a matter of seconds that took hours or which did not complete when built with PRIMUS or the Small Bonsai method alone.

Although we have presented an approach based on IBD segment overlaps for partitioning sets of distant relatives into their respective parental sides, relative to a focal individual or clade, it is likely that additional resolution could be gained by using IBD detected on sex chromosomes. At present, the Bonsai method uses only autosomal IBD to avoid considering the sexes of ancestral individuals along the paths connecting each pair of individuals when computing the likelihood of their relationship. Increased accuracy can also be obtained by using SNP-level information in our test of IBD overlap, such as opposite homozygotes, instead of IBD segments, as overlaps often occur between segments that are too short to be identified by existing IBD methods.

There is also potential to improve close relationship estimates by using phasing information. Williams et al. (2020) have demonstrated that half-sibling, avuncular, and grandparental relationships, which have been difficult to differentiate historically due to the fact that the total amount of IBD is the same for each of these relationship types, can be differentiated by making use of long-range phasing information. Phased IBD estimates, obtained from programs such as the PhasedIBD method of Freyman et al. (2020), could provide a considerable boost in accuracy for close relationships. Improved close relationships would lead to improved distant relationships due to the fact that the small pedigree structures being connected would be more accurate. The PhasedIBD method of Freyman et al. could also improve distant relationship estimates through more accurate inference of short IBD segments.

Although our method for detecting background IBD is able to distinguish background IBD from true IBD when the level of background IBD is low, the approach struggles when there is a significant quantity of background IBD. In such cases, other approaches for accounting for background IBD when inferring relationships can be used. One approach is to detect the amount of “self” IBD shared between homologous chromosomes in each individual in a pedigree. Assuming that all individuals in the pedigree come from the same population, the amount of self IBD provides an expected level of background IBD sharing between two haplotypes that can then be subtracted from each pairwise relationship of which the individual is a member. We find that this approach improves pedigree inference accuracy in practice.

The approach of using self IBD to adjust pairwise IBD estimates can also be an effective approach when inferring pedigrees with recent consanguinity. The current Bonsai method assumes that pedigrees are graphs without cycles. However, it is possible to include cycles when adding new individuals to the pedigree if individuals have substantial self IBD and their relationships with others are indicative of recent consanguinity. This approach can be used together with distributions that are specifically trained on relationships with consanguinity.

Approaches for inferring pedigrees in the context of background IBD and consanguinity are important for improving pedigree inference in all human populations. Although the theoretically maximal accuracy with which a pedigree can be inferred differs across human populations due to differences in demographic histories, it is likely that improvements in accuracy can be attained for all populations through improved methodology, such as the improvement of pairwise relationship inference by methods such as deep-learning trained in specific populations, the inclusion of additional consanguineous relationship types, and the inclusion of additional genetic information from sex chromosomes and mitochondrial DNA. By nature, pedigree inference is a complicated problem requiring methods that can handle a wide variety of pedigree structures and input data. However, our results show that the inference of large and sparse human pedigrees is tractable, and that accuracy will continue to increase as pedigrees become increasingly densely sampled.

## 6. Appendix

### 6.1. The probability of a pattern of IBD

Consider the induced subtree in a pedigree relating a set of genotyped individuals. This tree is shown with dashed red lines in Figure 4 with nodes of the tree indicated with red dots. Let *a*(*i*) denote the direct ancestral node of node *i* in this tree. For example, in the tree in Figure 4, we have *a*(1) = *A*_1_, *a*(6) = *A*_1_, *a*(2) = 6, *a*(3) = 6, *a*(4) = *A*_2_, *a*(5) = *A*_2_, *a*(*A*_1_) = *G* and *a*(*A*_2_) = *G*.

Assuming that all IBD segments are observed, we have

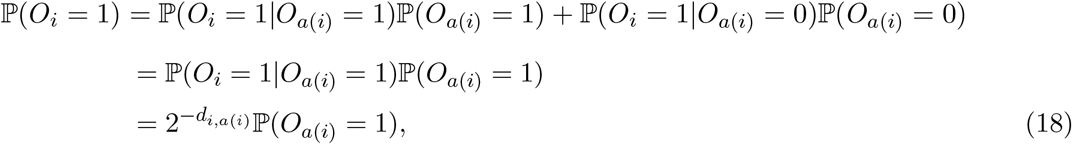

where *d*_*i,a*(*i*)_ is the number of meioses separating individual *i* from their ancestor *a*(*i*). Similarly, we have

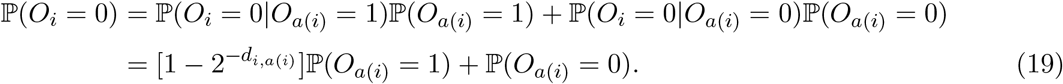

In the final lines of Equations (18) and (19), we have used the fact that the obability that an allelic copy is transmitted in one meiosis is 1*/*2.

Equations (18) and (19) establish a recursion for computing the probability of an observed presence and absence pattern from a given ancestral allelic copy at a single base of the genome. Defining

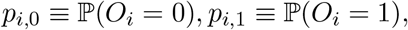

we can express the recursion compactly as

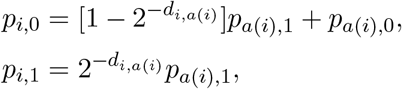

with the base conditions *p*_*g*,0_ = 0 and *p*_*g*,1_ = 1 for each chromatid, *g*, in *G*. The probability of an observed IBD sharing pattern {*O*_1_, …, *O*_*k*_} across *k* leaf nodes can be computed recursively using Equation (6).

### 6.2. Approximating the variance of *T*_1,2_

Here, we derive an approximation of the variance of the total length, *T*_1,2_, of IBD shared across the genotyped descendants of two acenstral individuals, *A*_1_ and *A*_2_. When we encounter a patch of IBD at a locus, the length of the patch can be approximated as the maximum length of | *𝒩*_1_| *×* | *𝒩*_2_| different IBD segments, where *𝒩*_*i*_ is the set of genotyped nodes below ancestor *A*_*i*_ at locus *m* in which the IBD segment is observed. This approximation comes from conceptualizing IBD sharing among the | *𝒩*_1_| IBD segment carrying descendants of *A*_1_ and the |*N*_2_| IBD segment carrying descendants of *A*_2_ as | *𝒩*_1_| *×* | *𝒩*_2_| independent segments with a single point at which all segments overlap. The length of the merged segment to one side of this focal point then has a distribution given by the maximum of | *𝒩*_1_| *×* | *𝒩*_2_| exponential random variables whose means depend on the degree of separation between the corresponding pairs of leaf individuals. To simplify matters, we assume that the length of the full merged overlapping segment (not just to the left or right) is exponentially distributed.

This approximation is an oversimplification of the IBD sharing pattern because the segments are not truly independent and need not overlap at a single point. Moreover, under this approximation, the length of the merged segment would be the maximum over sums of identically distributed random variables, representing the sum of the length of a segment to the right of the center point and the length of the segment to the left. However, we are not overly concerned with these drawbacks of the conceptualization because our main goal is to obtain an accurate, yet simple approximation of the variance of the distribution. We also assume that no member of 𝒩_*i*_ is the direct ancestor of another member of the set, which holds in practice if we drop all individuals from 𝒩_*i*_ who are descended from others.

The length, 𝓁_*i,j*_, of an IBD segment between leaf nodes *i* and *j* is can be modeled as an exponentially distributed random variable with mean length *µ*_*ij*_ = *L*_*genome*_*/d*_*i,j*_*R*, where *d*_*i,j*_ is the degree of relationship between them and *R* is the expected number of recombination events, genome wide, in one meiosis (Huff et al. 2011). When the length of the genome is expressed in centimorgans (cM), the expected number of recombination events in the genome is *L*_*genome*_*/*100. Thus, the expected length in cM of an IBD segment between individuals *i* and *j* separated by *d*_*i,j*_ meioses is *µ*_*ij*_ = 100*/d*_*i,j*_.

Let *L*_1,2_ denote a random variable describing the length of the segment formed by merging all segments at a given locus *m* between descendants of *A*_1_ and *A*_2_. If the lengths of all segments at this locus were independent, their merged length in our conceptualization would have a distribution given by the maximum over independent exponentially distributed random variables with means 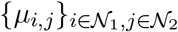.

If the leaf nodes with observed IBD at the marker are *𝒩*_1_ and *𝒩*_2_, then we have 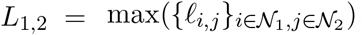. Under this condition, the cumulative density function (CDF) *F*_*L*_(𝓁; *𝒩*_1_, *𝒩*_2_) of *L* is

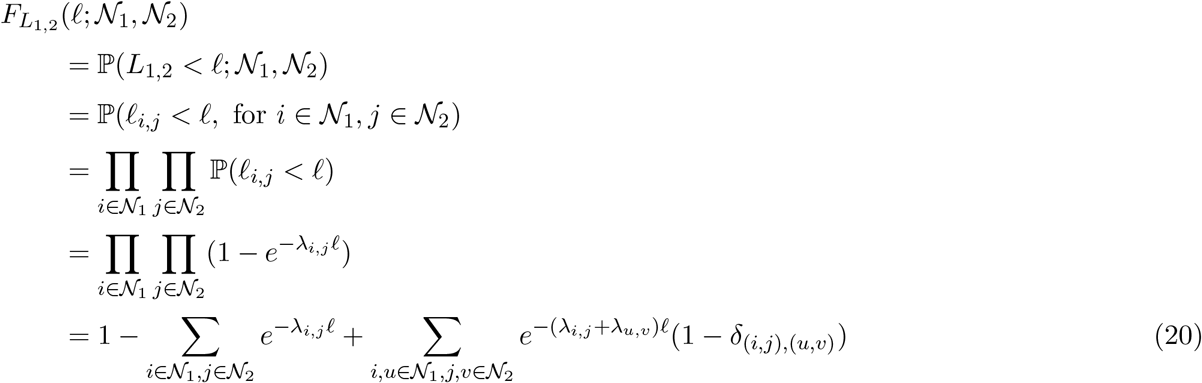

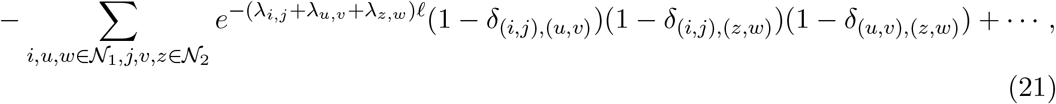

where *λ*_*i,j*_ = 1*/µ*_*i,j*_ = *d*_*i,j*_*/*100 and δ_(*a,b*),(*c,d*)_ is the Kronecker delta between tuples (*a, b*) and (*c, d*), which is equal to one when (*a, b*) = (*c, d*) and zero, otherwise.

The sets *𝒩*_1_ and *𝒩*_2_ are, themselves, random variables. Summing over all sets *𝒩*_1_ and *𝒩*_2_, we have

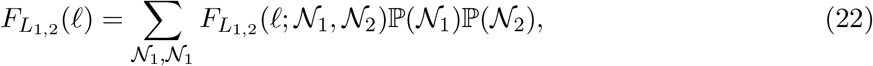

where the probabilities 𝕡 (*𝒩*_1_) and 𝕡 (*𝒩*_2_) are probabilities of observing IBD in the sets of leaf nodes below *A*_1_ and *A*_2_ and can be approximated using the recursion in Equation (6).

Over the length of the genome, the number *𝒩*_1,2_ of IBD segments between the descendants of *A*_1_ and *A*_2_ is approximately Poisson distributed with mean (1 *− 𝕡* (*ℐ*^*c*^)^2|*G*|^)*L*_*genome*_*/E*[*L*_1,2_]. This rate comes from the fact that the average total amount of the genome in a patch of IBD is (1 *−* P(*ℐ*^*c*^)^2|*G*|^)*L*_*genome*_ while the average length of any given segment is *E*[*L*_1,2_]. When the lengths of IBD are short and far apart, which they are when the degree between *A*_1_ and *A*_2_ is large, this is a reasonable approximation. This is precisely the regime in which the distribution in Equation (15) is most useful.

The total length *T*_1,2_ of merged IBD among the descendants of *A*_1_ and *A*_2_ is

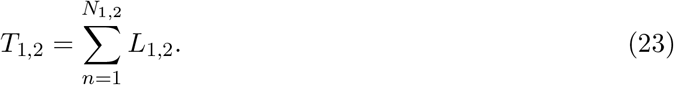

We can derive the variance of *T*_1,2_ using the law of total variance as

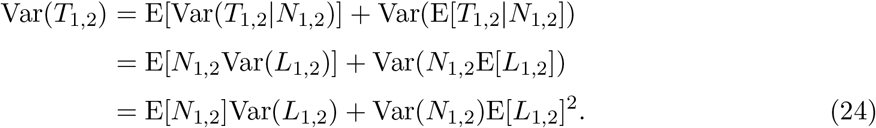

Note that because *N*_1,2 ∼_ Poisson ((1 − ℙ(ℙ^c^)^2 ℙGℙ^) *L*_genome_/E[*L*_1,2}_].

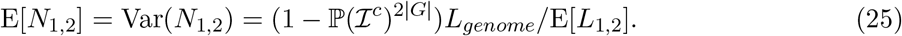

So Equation (24) simplifies to

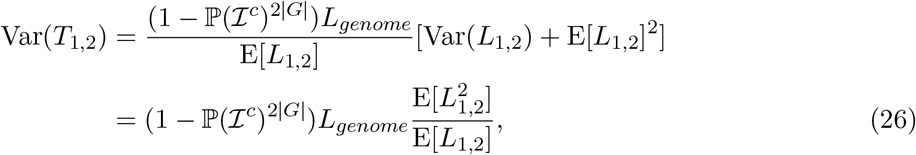

where we have used the fact that Var(*X*) = E[*X*^2^] − E[*X*]^2^.

It remains to find E[*L*_1,2_] and 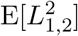. Using the CDF of *L*_1,2_ in Equation (22) and the fact that E[*X*^*m*^] = *m*! ∫_ℝ_ *x*^*m*−1^[1 − *F*_*X*_(*x*)]*dx*, we have

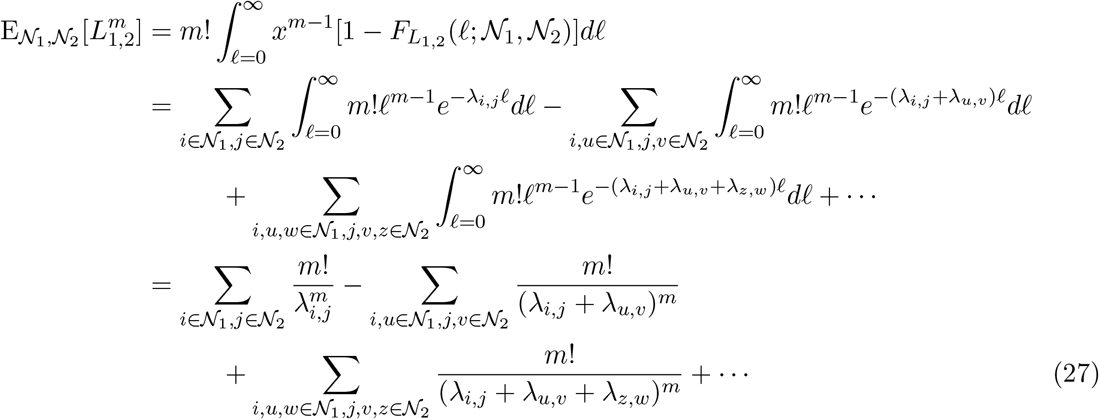

where the integrals in Equation (27) can be evaluated by noting that they are essentially expressions for the moments of exponential random variables with parameters *λ*_*i*_, (*λ*_*i*_ + *λ*_*j*_), (*λ*_*i*_ + *λ*_*j*_ + *λ*_*k*_), etc.

Thus, we can use Equation (27) to compute

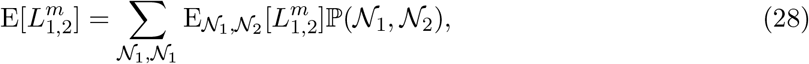

where ℙ (*𝒩*_1_, *𝒩*_2_) is the probability of observing IBD segments at the leaves *𝒩*_1_ and *𝒩*_2_, and is approximated using the recursion in Equation (6). We then plug Equation (28) in to obtain the variance of *T*_1,2_ in Equation (26).

In practice, it is too computationally demanding to compute the sums in Equation (28) because the terms 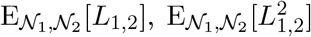, and 𝕡 (*𝒩*_1_, *𝒩*_2_) are not fast to compute in large quantities. However, the probabilities 𝕡 (*𝒩*_1_, *𝒩*_2_) can be computed quickly enough, allowing us to find the most likely sets of leaf nodes, 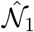 and 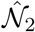, with observed IBD. Thus, in practice we use an approximation in which we assume that the most likely IBD pattern has been observed and we compute

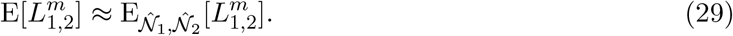

The assumption used in this approximation is that most patterns of observed IBD at the leaves are unlikely compared with the most likely patterns and that most likely patterns of IBD will yield similar moments 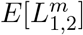.

### 6.3. Re-rooting the DRUID estimator

In some scenarios, the common ancestors, *A*_1_ and *A*_2_, of sets of individuals *𝒩*_1_ and *𝒩*_2_, may not be related through a common ancestor or ancestral pair of both *A*_1_ and *A*_2_. In particular *A*_2_ can be the direct descendant of *A*_1_, or vice versa. This scenario, along with the scenario treated in Section 3.5.3 in which *𝒩*_1_ and *𝒩*_2_ are connected through their common ancestors, covers all possible ways in which *𝒩*_1_ and *𝒩*_2_ can be connected such that they are mutually related, i.e., so that they share a common ancestor.

We now describe an approach for computing the generalized DRUID estimate when *A*_2_ is descended from an individual *A* who is the common ancestor of only a subset of *𝒩*_1_. We consider *A* to be any node ancestral to some node in *𝒩*_1_, including any member of *𝒩*_1_ itself.

Let Λ_1_(*A*_1_) denote the induced subtree in pedigree *𝒫*_1_ that relates *A*_1_ and their descendants *𝒩*_1_. To obtain the generalized DRUID estimate when *A*_2_ is descended from *A*, we re-root the tree Λ_1_(*A*_1_) at *A* to obtain a re-rooted tree 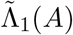. We then compute the generalized DRUID estimate from Section 3.5.3 using the re-rooted tree 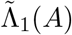. The estimate between *A* and *A*_2_ obtained using Equation (9) applied to 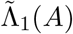 and Λ_2_(*A*_2_) is then the number of meioses separating *A* and *A*_2_.

The one complication is that *A*_2_ can be descended from both *A* and a spouse *A*^*’*^, who is also an ancestor of one or more of *A*’s genotyped descendants *𝒩*_*A*_. In this case, *A*_2_ is more closely related to *𝒩*_*A*_ than to *𝒩*_1_ \ *𝒩*_*A*_ by one degree. We solve this problem by representing the clades of shared descendants *twice* on the re-rooted tree, obtaining a multi-labeled tree (Figure 15). In contrast, if *A*_2_ is descended from *A* and a spouse *A*^*“*^ who is not ancestral to any genotyped descendants, we do not duplicate the descendants of *A* on the tree.

**Figure 15.**
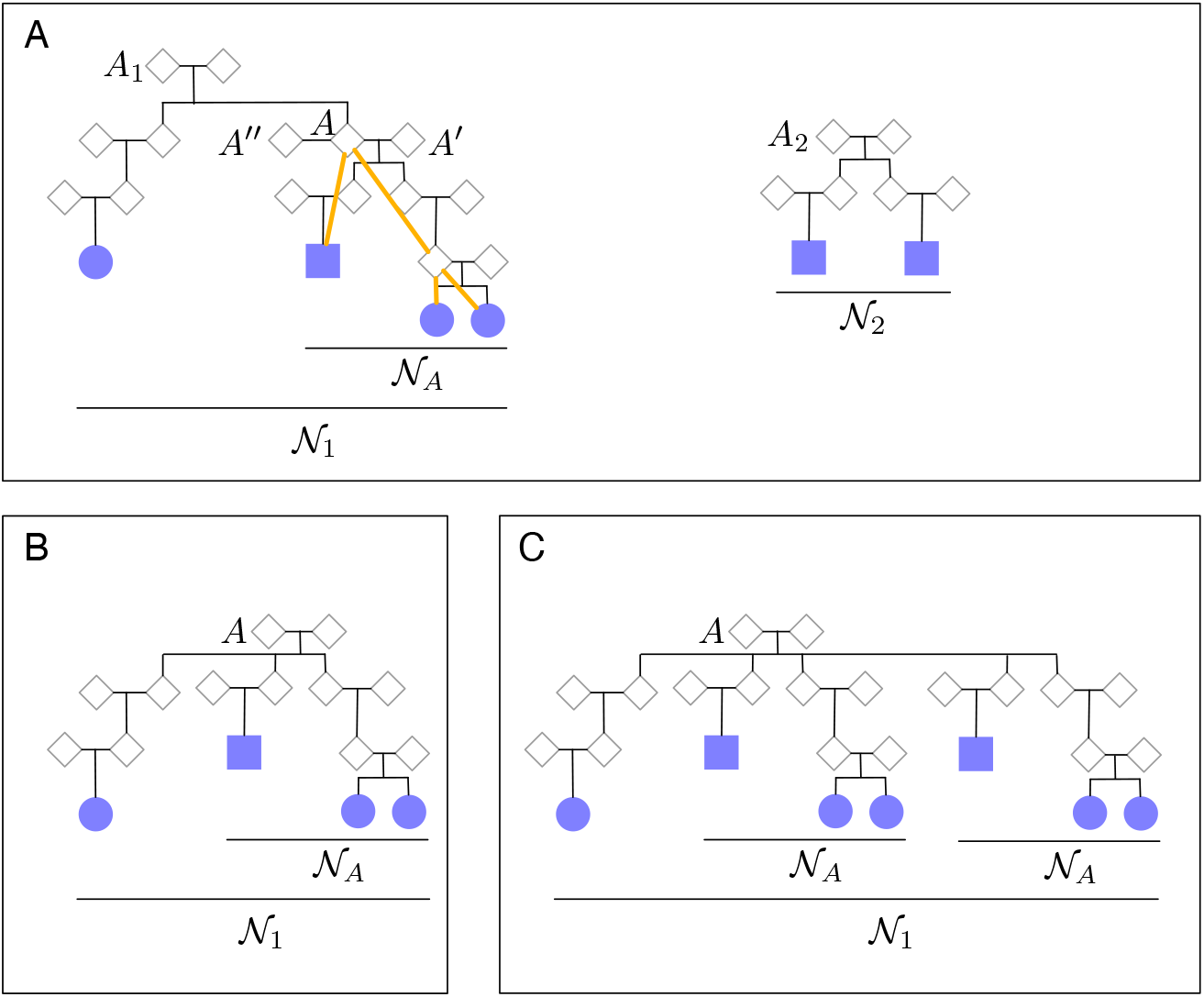
Re-rooting the DRUID estimator. (A) The pedigrees relating *𝒩*_1_ and *𝒩*_2_, respectively. The induced subtree relating the descendants of internal node *A* is shown in orange. (B) The re-rooted tree when *A*_2_ is descended from *A* and *A*^*“*^.(C) The re-rooted tree when *A*_2_ is descended from *A* and *A*^*’*^.

## 7. Acknowledgements

We would like to thank the employees and research participants of 23andMe who made this research possible. We would also like to thank Amy Williams and Ying Qiao for helpful discussions. Members of the 23andMe Research Team are Michelle Agee, Stella Aslibekyan, Elizabeth Babalola, Robert K. Bell, Jessica Bielenberg, Katarzyna Bryc, Emily Bullis, Briana Cameron, Daniella Coker, Gabriel Cuellar Partida, Devika Dhamija, Sayantan Das, Sarah L. Elson, Teresa Filshtein, Kipper Fletez-Brant, Pierre Fontanillas, Pooja M. Gandhi, Karl Heilbron, Barry Hicks, David A. Hinds, Karen E. Huber, Yunxuan Jiang, Aaron Kleinman, Katelyn Kukar, Keng-Han Lin, Maya Lowe, Marie K. Luff, Jennifer C. McCreight, Matthew H. McIntyre, Steven J. Micheletti, Meghan E. Moreno, Joanna L. Mountain, Sahar V. Moza_ari, Priyanka Nandakumar, Elizabeth S. Noblin, Jared O’Connell, Aaron A. Petrakovitz, G. David Poznik, Anjali J. Shastri, Janie F. Shelton, Jingchunzi Shi, Suyash Shringarpure, Chao Tian, Vinh Tran, Joyce Y. Tung, Xin Wang, Wei Wang, Catherine H. Weldon, and Peter Wilton.

## 8. Declarations of interests

The authors are employees of 23andMe, Inc., and hold stock or stock options in 23andMe.

## 9. Web resources

The Bonsai code is available at https://github.com/23andMe/bonsaitree.

## 10. Supplemental figures

**Figure S1.**
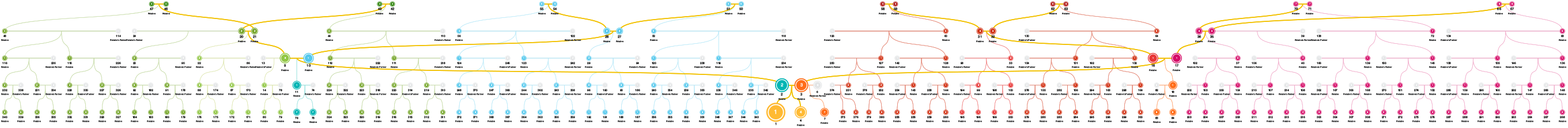
Example of a large pedigree for testing the Big Bonsai method. 100 such pedigrees were simulated using the approach described in Section 3.6.5. The focal individual is labeled “1.”

**Figure S2.**
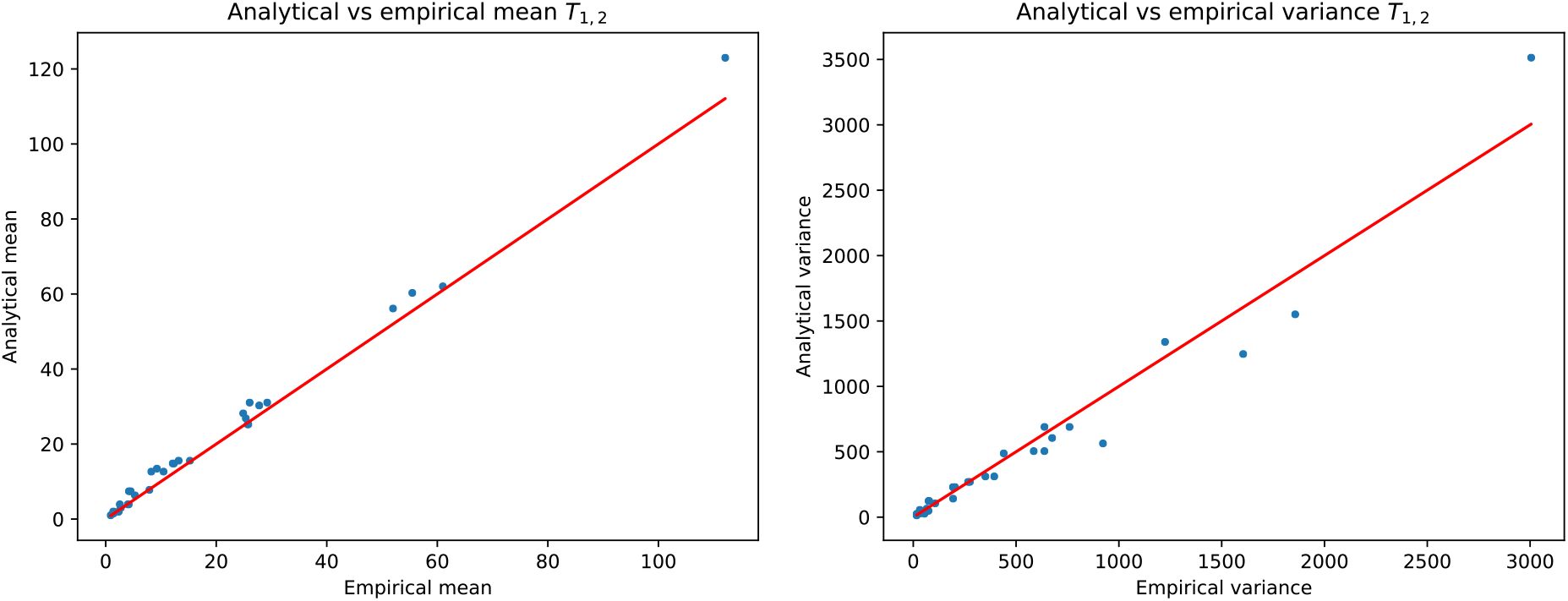
Analytical versus empirical mean and variance of *T*_1,2_. Analytical means were computed using Equation (12) and analytical variances were computed using Equation (13). Empirical means and variances were computed using simulated pedigrees comprised of two small pedigrees *𝒫*_1_ and *𝒫*_2_ connected through either one or two common ancestors. Pedigree_1_ had a randomly generated structure simulated by starting with the pair of root individuals and their two children. At each subsequent generation, each leaf node had a probability 1*/*2 of having one offspring and 1*/*2 of having two offspring. Pedigree *𝒫*_1_ was extended down from the root nodes until the total number of leaves was |*𝒩*_1_|. Pedigree *𝒫*_2_ was simulated in the same way, but independently of *𝒫*_1_. One common ancestor *A*_1_ of *𝒫*_1_ was then connected to one common ancestor *A*_2_ of *𝒫*_2_ through either one or two common ancestors, *G*. The degrees 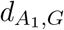 and 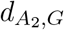 were set to either 3 or 5 and the number of common ancestors |*G*| was either 1 or 2. Thus, the genealogical degrees between *A*_1_ and *A*_2_ were in the set {5, 6, 7, 8, 9, 10}. The number of leaves in each pedigree was either *𝒩*_*i*_ = 2 or *𝒩*_*i*_ = 4. We ran 10 simulation replicates for each configuration of 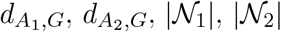, and |*G*|.

**Figure S3.**
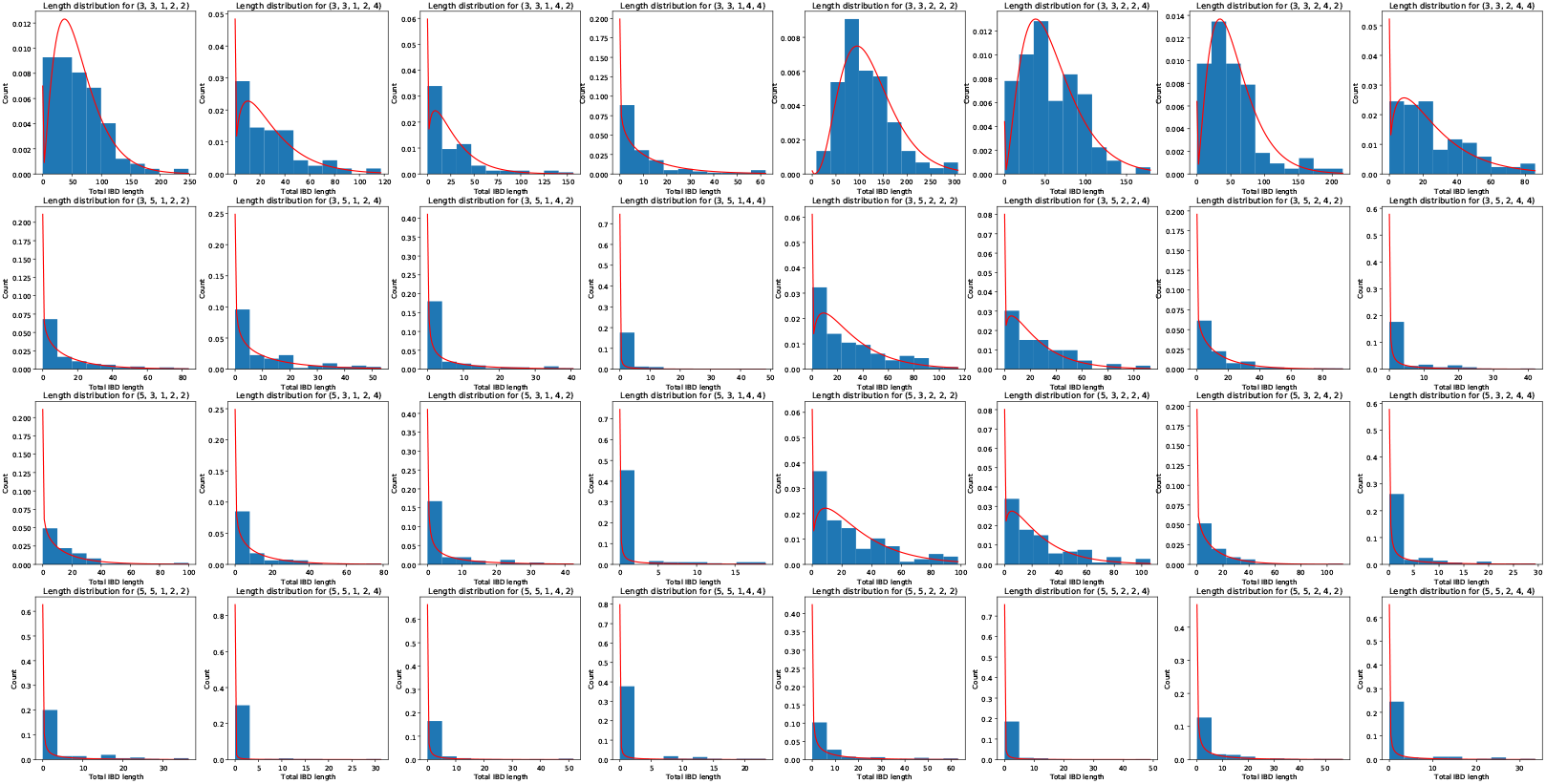
Analytical versus empirical distributions of *T*_1,2_. Analytical distributions were computed using Equation (15). Empirical distributions were simulated in the manner described in Figure S2.

## Notes

https://github.com/23andMe/bonsaitree

